# Molecular Bonsai: Elucidating the design principles for engineering plant organ size

**DOI:** 10.1101/2025.06.22.660808

**Authors:** Tawni Bull, Jared Van Blair, Sam Sutton, Hadley Colwell, Arjun Khakhar

## Abstract

Enhancements to crop morphology, such as the semi-dwarfing that helped drive the green revolution, are often driven by changes in gene expression. These are challenging to translate across species, which slows the rate of crop improvement. Synthetic transcription factors (SynTFs) offer a rapid alternative to generate targeted alterations to gene expression. However, the complexity of developmental pathways makes it unclear how to best apply them to predictably engineer morphology. In this work, we explore whether mathematical modeling can guide SynTF-based gene expression modulation to help elucidate the design principles of engineering organ size. We targeted genes in the phytohormone, gibberellin (GA), signaling pathway, which is a central regulator of cell expansion. We demonstrate that modulation of GA signaling gene expression can generate consistent dwarfing across tissues and environments in *Arabidopsis thaliana*, and that the degree of dwarfing is dependent on the strength of regulation, as predicted by modeling. We further validate the model’s predictive power by demonstrating its capacity to predict the qualitative impacts of different regulatory architectures for engineering organ size. Additionally, we develop expression parameterized models to quantitatively predict organ size and elucidate how temperature will affect growth. Finally, we show that these insights can be generalized for engineering organ size in tomato (*Solanum lycopersicum*). This work creates a framework for predictable engineering of an agriculturally important trait across tissues and plant species. It also serves as a proof-of-concept for how mathematical models can guide SynTF-based alterations in gene expression to enable bottom-up design of plant phenotypes.

**Significance Statement:** While traditional breeding approaches have identified mutations that enhance crop performance via targeted gene expression changes, these are not easily translated across varieties and species. Synthetic transcription factors (SynTFs) offer an avenue to generate such changes *de novo*, but the optimal regulatory architectures necessary to generate desired phenotypes remain unclear. We demonstrate how mathematical models can be used to guide SynTF deployment and elucidate the design principles for engineering organ size, an agriculturally important trait, via modulation of gibberellin signaling. In addition to revealing regulatory architectures that can consistently increase or decrease organ size across a range of tissues, environments, and plant species, this work demonstrates how model-guided SynTF-based modulation of gene dosage can be used to predictably engineer plants.

## Introduction

Morphological alterations, such as changes to organ size, make up a large fraction of crop trait improvements that have been selected over the course of domestication (1). For example, semi-dwarfed stems in cereals dramatically increased yields during the Green Revolution by both enhancing resilience to wind damage, as well as improving productivity per unit area by enabling higher density planting (2). Studies into the genetic basis of these improved morphologies reveal that the majority are generated by *cis*-regulatory changes that alter gene expression (3–10). However, translating these insights to optimize traits, or introducing them into new varieties or species using traditional breeding strategies is challenging due to epistasis, cross regulation, and incomplete understanding of plant *cis* regulation. This necessitates trait optimization in each new genetic background, which significantly slows crop enhancement. There is an urgent need to accelerate this process to address the twin challenges posed by climate change and increasing demand for future food security.

The bottom-up generation of gene expression changes using synthetic control systems represents a faster alternative for crop improvement (**Fig. 1A**). Cas9-based synthetic transcription factors (SynTFs) have been demonstrated to enable layering of additional regulation on top of native signaling at endogenous loci in plants (11, 12). They are also orthogonal to native developmental pathways, reducing challenges to translation across genetic backgrounds and species. This makes them an ideal tool for modulating the expression of genes to engineer developmental phenotypes, such as organ size. However, design principles for organ size engineering with SynTFs remain opaque due to many open questions. These include: 1) which gene(s) can be modulated to generate robust and predictable changes in organ size, 2) what is the dose-dependency between gene expression and phenotype, and 3) Can utilizing SynTFs both increase and decrease organ size? It is also unclear whether this approach would generate consistent phenotypic changes across tissues, environments, and species.

**Figure 1.**
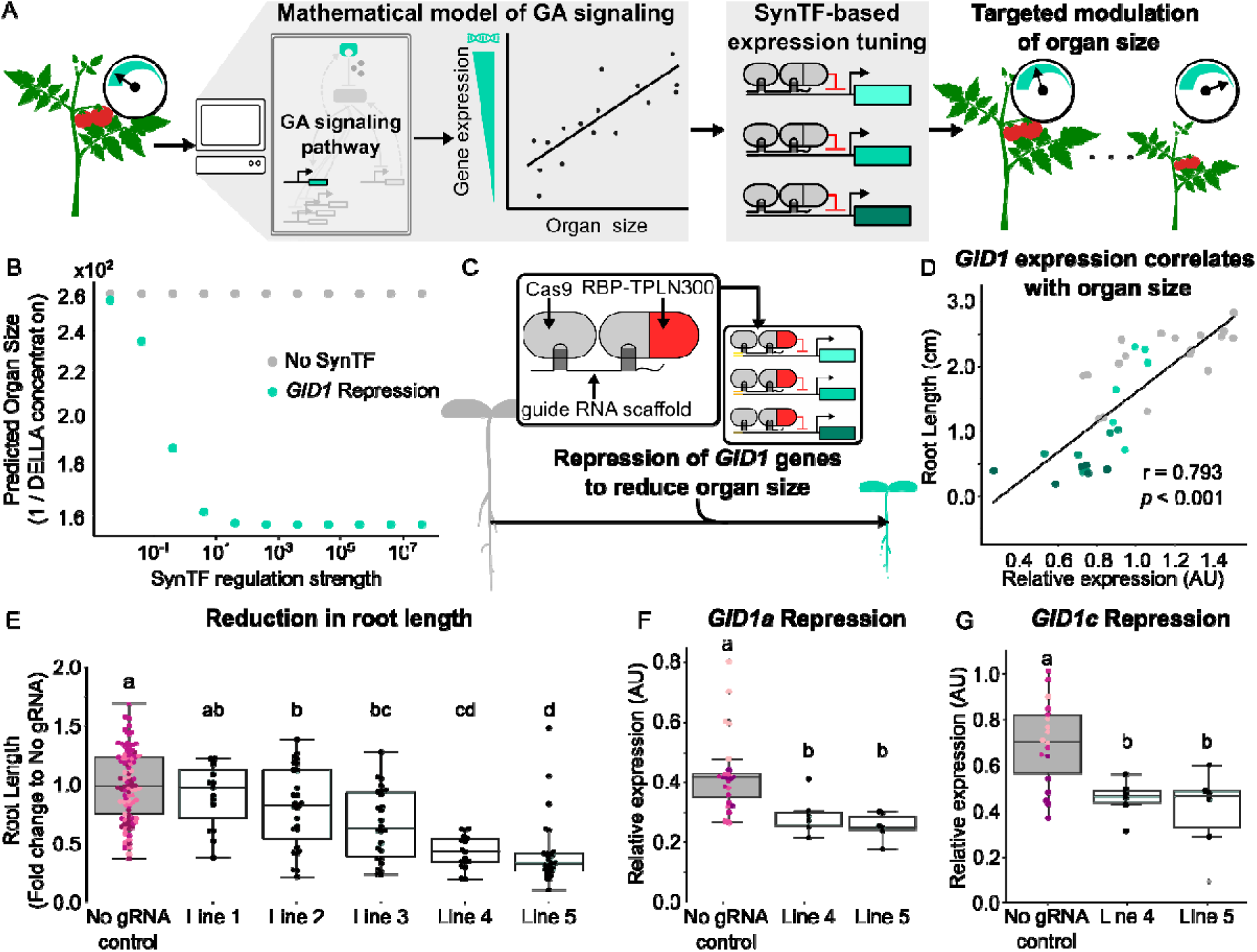
SynTF-based repression of GID1 generates dose-dependent dwarfing predicted via modeling in. A. thaliana. A) Schematic depicting the use of mathematical modeling to guide SynTF-based modulation of gibberellin signaling gene expression and to elucidate the design principles for predictably engineering organ size. B) Plot summarizing the predicted reciprocal of DELLA protein concentration at steady state, used as a proxy for organ size, from simulations of the GA signaling network with either No SynTF (gray) or a repressor targeted to regulate *GID1* (teal), across a range of different SynTF regulation strengths. C) A schematic of the Cas9-based SynTF used to implement repression on the three different *GID1* paralogs in *A. thaliana.* D) Scatterplot that demonstrates the correlation between root length and the sum of normalized *GID1a* and *GID1c* expression. Every dot of the same color corresponds to an independent biological replicate of either the no gRNA control lines (gray) or *GID1* repression lines 1, 4, and 5 (progressively darker shades of teal, respectively). E) Boxplots summarizing the fold change in root length of plant lines with SynTF-based repression of *GID1* (teal) compared to a population of no gRNA control plant lines (gray, n=4). F, G) Boxplots summarizing the expression of *GID1a* (F) and *GID1c* (G) normalized by housekeeping gene expression (*PP2A*) in seedlings of a population of either no gRNA control lines (gray, n=3) or the *GID1* repression lines with the strongest phenotypes (teal). Every dot of the same color within each boxplot corresponds to an independent biological replicate of the same genotype. Different letters represent statistically significant differences (One-way ANOVA followed by Tukey HSD test, *p* < 0.05).

Prior studies have revealed that the signaling pathway of the phytohormone gibberellin (GA) is a core determinant of cell expansion, and hence organ size, across plant species (13–16). GA associates with its receptor, GID1 (GIBBERELLIN INSENSITIVE DWARF 1), which triggers association of GID1 and a family of transcriptional modulators, called DELLA proteins, leading to their ubiquitination and subsequent proteasomal degradation (17–19, **Fig. S1A**). These DELLA proteins function as negative regulators of cell expansion and, therefore, increases in GA tend to increase organ size (17, 20, 21). Additionally, it has been shown that DELLAs are also responsible for implementing multiple transcriptional feedback loops that maintain GA homeostasis via repression of their own expression and activation of both *GID1* and GA biosynthesis gene (GA20 oxidases, GA3 oxidases) expression (22, **Fig. S1A**). This then generates post-translational feedback via modulation of the concentration of GA and the sensitivity of its perception by *GID1*, which ultimately alters the rate of DELLA protein degradation.

These multi-layered transcriptional and post-translational feedback loops make it challenging to intuitively predict how changing the expression of a gene in this network with SynTFs would alter DELLA protein levels, and hence organ size. For example, increasing *GID1* expression would decrease DELLA protein levels initially by increasing GA-induced DELLA degradation. However, because of feedback, the reduction in DELLA protein would result in an increase in *DELLA* transcription and a decrease in *GA20ox* transcription resulting in less GA biosynthesis, both of which would increase DELLA protein levels. Thus, the final impact on DELLA protein concentration is not obvious. A mathematical model of this feedback network parameterized using data from roots of *Arabidopsis thaliana* GA signaling mutants (22) provides an avenue to overcome this issue via simulation of the network dynamics to a steady state. Additionally, a modified version of this model that incorporates SynTF-based modulation of GA signaling gene expression, along with the native feedback regulation, could enable prediction of steady state DELLA protein levels, and hence organ size, in engineered plants.

Another challenge includes that the model’s predictive power remains unexplored in tissues besides roots and in other plant species. Existing studies that use such models to guide modulation of GA regulated growth have focused on GA biosynthesis genes in roots (9, 11, 23, 24). Additionally, studies of the role of GA signaling genes, namely *GID1* and *DELLA*, in controlling organ size have largely focused on knockout mutants or transgene overexpression. These approaches create extreme expression changes as well as disrupt spatiotemporal expression patterns and native feedback signaling, making it challenging to use their observations to interrogate the impact of gene expression on organ size (6, 17, 22, 25–32). Thus, it remains unknown whether the expression of GA signaling genes can be modulated to levels within these extremes to predictably engineer organ size.

In this work we seek to elucidate the design principles for SynTF-based organ size engineering via modulation of GA signaling gene expression. Specifically, we explore whether mathematical models can elucidate regulatory architectures that generate robust and predictable changes in organ size (**Fig. 1A**). This includes comparing the relative efficacy of regulatory architectures for both decreasing and increasing organ size, as well as characterizing the relationship between regulation strength and phenotype across multiple transgenic lines and in varying environmental conditions. We also test whether these circuits generate organ size changes across different tissues, and if the phenotypes they generate are robust to environmental fluctuations. Furthermore, we collected gene expression measurements from these transgenic lines to develop expression parameterized models that can quantitatively predict organ size. Finally, we test whether the insights gained from the model in *A. thaliana* can be generalized to engineer organ size in an agriculturally important crop, *Solanum lycopersicum* (tomato).

## Results

### Model-guided repression of *GID1* generates predicted dwarfing in roots

We implemented a modified version of a published ordinary differential equation-based model of the GA signaling network (22) in python to simulate the impact of SynTF-based repression of *GID1* expression on DELLA protein levels in the context of DELLA-mediated transcriptional feedback (**Fig. S1**). The original model’s equations simulate the transcriptional and post-translational feedback loops inherent in the GA signaling pathway. Additional equations were added to the model to simulate SynTF transcription and translation, and repression of *GID1* by the SynTF was modeled using a Hill equation, as described in the methods section (**Fig. S1**). Because DELLA has been shown to inhibit cell expansion via regulation of downstream transcription factors (TFs) (33), we used the reciprocal of DELLA protein concentration as a qualitative metric of predicted organ size. We hypothesize that this kind of linear relationship between DELLA concentration and organ size would likely not apply to very high DELLA concentrations, where the TFs that it regulates are saturated, or very low concentrations, where DELLAs would preferentially interact with the subset of TFs that have the strongest interaction strengths. However, we reasoned that this assumption should be reasonable for the moderate dosage differences from wildtype levels typically generated by SynTFs (11, 34). Our simulations predicted that the repression of *GID1* expression would reduce organ size via increasing DELLA protein concentrations. They also predicted that the strength of repression of *GID1* expression would be correlated with the reduction in organ size over a range of regulation strengths (**Fig. 1B**). Based on these predictions, we built *A. thaliana* transgenic lines with SynTFs targeted to regulate the three paralogs of the *GID1* family, *GID1a* (AT3G05120), *GID1b* (AT3G63010) and *GID1c* (AT5G27320).

We utilized a SynTF design that enables flexible implementation of activation and repression across multiple genes *in planta* (11). It consists of a constitutively expressed nuclease active Cas9 and RNA-binding proteins, PCP and MCP, fused to previously validated transcriptional effector domains, specifically truncations of the TOPLESS repressor for repression and the DREB2A trans-activator for activation (**Fig. 1C**). When these components are co-expressed with a guide RNA scaffold (gRNA), assembly of the desired transcription factor occurs at the promoter of the target gene (11). The gRNAs have a truncated target site, which programs Cas9 to bind rather than cut DNA (35), and motifs at their 3’ end to recruit the appropriate RNA-binding protein-transcriptional effector fusion (36). Multiple transgenic *A. thaliana* lines expressing the SynTF components and gRNAs that assemble a repressor at the promoters of each of the *GID1* genes, simultaneously, were generated (11). Additionally, we generated *A. thaliana* lines that expressed all the SynTF components but no gRNAs as controls to account for the impact of selection and SynTF expression on phenotype.

Seedlings of both *GID1* repression (SynTF repressor targeted to each *GID1*) and no gRNA control lines were grown in tandem on MS agar plates for eight days. We subsequently measured primary root length as this was the tissue for which the original model was parameterized. Four out of five *GID1* repression lines assessed showed significant reduction in root length when compared to a population of no gRNA control lines (**Fig. 1E**). We observed a range of root dwarfing levels in these lines, spanning a 0.41-to-0.81-fold change in mean root length compared to the no gRNA control population. Based on our simulations, we hypothesized that this variation in phenotype strength was due to the expected variation in the strength of *GID1* repression across independent transgenic events associated with random insertion and copy number variation of SynTF components.

To test this hypothesis, we conducted RT-qPCR to measure the degree of *GID1* repression across these lines. We observed significant *GID1a* and *GID1c* repression across lines compared to the no gRNA controls (**Fig. 1F, G**). However, no *GID1b* repression was observed in any line (**Fig. S2A**). We hypothesize that this is due to an inefficient gRNA target sequence. We then measured the correlation between tandem measurements of *GID1* expression and root length and observed significant positive correlations for *GID1a* (r = 0.745, *p* < 0.001) and *GID1c* (r = 0.75, *p* < 0.001) (**Fig. S2C, D**). We observe an even stronger correlation (r = 0.793, *p* < 0.001) between the total *GID1a* and *GID1c* expression in the plant and root length, which is consistent with these genes being functionally redundant (**Fig. 1D**). These results are consistent with model predictions showing that increasing SynTF repression of *GID1* would increase DELLA protein levels, leading to larger reductions in organ size (**Fig. 1B**). We also noted that the *GID1* repression was most visible in tissues sampled at night, but non-significant in tissues collected in the day (**Fig. S2B**). This is consistent with the reported circadian oscillation of *GID1* expression, with low expression in the day making repression challenging to observe and high expression at night revealing the impact of SynTF-based repression (37). Taken together, these results validate *GID1* as a good target for SynTF-based engineering of root dwarfing.

### SynTF-based repression of *GID1* results in smaller organ size across multiple tissues

As the parameterization of our model is based on data collected from *A. thaliana* seedling roots, it was possible that this circuit would not generate the same phenotype across other tissues and developmental times (22). To explore this, we measured shoot phenotypes at different developmental times (**Fig. 2**), including hypocotyl length and leaf size, in a subset of the transgenic lines with SynTF-based repression of *GID1* described above.

**Figure 2.**
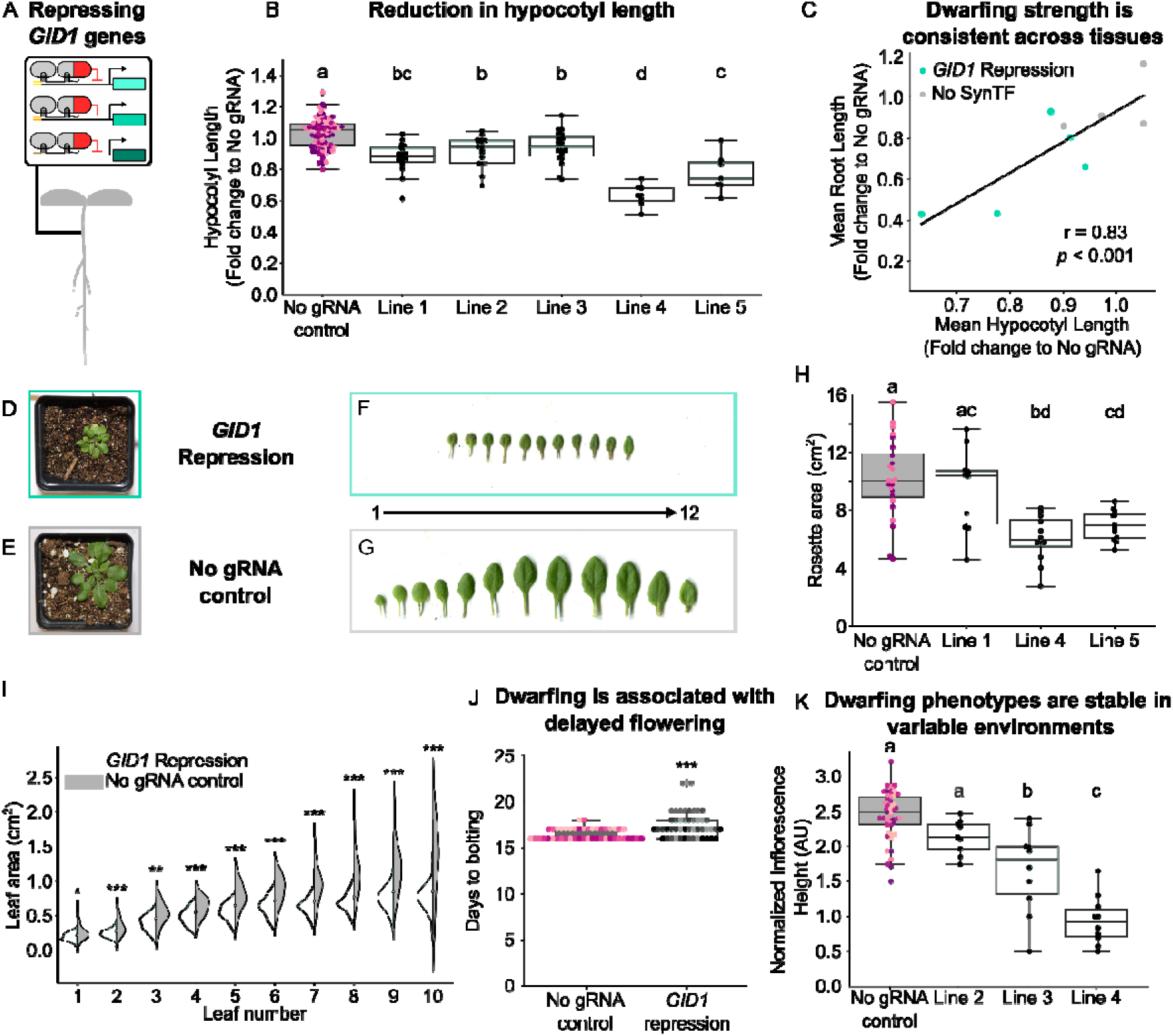
SynTF-based GID1 repression generates consistent dwarfing across tissues and environments. A) Schematic depicting the genetic circuit enabling SynTF-based repression of *GID1* genes. B) Boxplots summarizing the fold change in hypocotyl length of the *GID1* repression lines (teal) compared to a population of no gRNA control plant lines (gray, n=5). C) Scatterplot that depicts the Pearson’s correlation between mean root length and mean hypocotyl length of *GID1* repression lines (teal) and no gRNA control lines (gray). D,E,F,G) Representative pictures of rosettes (D,E) and dissected leaves arranged according to developmental age (F,G) from *GID1* repression (teal), and no gRNA control lines (gray). H) Boxplots summarizing the total rosette area of three plant lines with SynTF-based repression of *GID1* (teal) compared to a population of no gRNA control plant lines (gray, n=2). I) Stacked violin plot showing the distribution of leaf areas for plant lines with SynTF-based repression of *GID1* (teal, 3 lines with 10 biological replicates) compared to the no gRNA control plant lines (gray, 2 lines with 10 biological replicates) for each leaf in developmental time where leaf 10 is youngest measured. J) Boxplots summarizing the days to bolting of the *GID1* repression lines (teal, n=3) compared to the population of no gRNA control lines (gray, n=5). K) Boxplots summarizing the height of inflorescences normalized by the number of days post bolting of plant lines with SynTF-based repression of *GID1* (teal) compared to a population of no gRNA control plant lines (gray, n=5) grown in a variable greenhouse environment. Every dot of the same color within each boxplot corresponds to an independent biological replicate of the same genotype. Different letters represent statistically significant differences (One-way ANOVA followed by Tukey HSD test, *p* < 0.05). Asterisks represent results of a Welch’s two sample *t*-test (*p* < 0.05), * corresponds to *p* < 0.05, ** corresponds to *p* < 0.005, and *** corresponds to *p* < 0.0005.

Prior work has demonstrated that the GA signaling pathway plays a major role in hypocotyl elongation (3, 38, 39). Therefore, if the model predicted an increase in DELLA concentration upon *GID1* repression, we would expect to observe a decreased ability for hypocotyl elongation. To test this hypothesis, we instigated hypocotyl etiolation in newly germinated seedlings of *GID1* repression lines (**Fig. 2A**) in tandem with no gRNA controls. We observed that the *GID1* repression lines showed a significant decrease in hypocotyl length when compared to the no gRNA control population (**Fig. 2B**). Interestingly, this included the line that did not show significant reductions in primary root length. This is likely due to a smaller variability in hypocotyl phenotypes (CV = 14.9%) when compared to the root phenotypes (CV = 45.2%) we observed across all lines. We also observed a strong correlation (r = 0.83, *p* < 0.001) between the strength of dwarfing, quantified as the fold change in organ size compared to the no gRNA controls, observed in roots and hypocotyls across these lines (**Fig. 2C**). These observations further validate the role played by GA signaling in hypocotyl elongation (3, 38) and demonstrate the capacity of SynTF-based *GID1* repression to generate consistent dwarfing across different tissues.

To further explore our SynTF’s capacity to generate dwarfing across tissues, we also measured individual leaf area and calculated total rosette size in *GID1* repression lines and observed a significant decrease in both phenotypes compared to the no gRNA control population (**Fig. 2D, E, H**). Like our measurements of hypocotyl length, lines with the strongest dwarfing in roots had the largest changes in rosette size (**Fig. 2H**). These results, in conjunction with the correlation between *GID1* expression and root length (**Fig. 1D**), show that repressing *GID1* expression generates a consistent degree of dwarfing across tissues that is dependent on the strength of SynTF regulation generated from T-DNA insertional variation.

Interestingly, we observed that in the distribution of leaf areas across developmental time, the lower bound of the distribution is not decreased in the *GID1* repression lines when compared to the control but the upper bound is, driving an overall decrease in leaf area (**Fig. 2F, G, I**). These observations are consistent with the previously reported phenotypes of *GID1* knockouts (17) and with our hypothesized reduction in cell expansion in the *GID1* repression lines compared to the no gRNA control lines. The invariant lower bound in leaf size is due to processes inherent to leaf development wherein cells transition from a phase of mainly cell division, where lines behave similarly, to cell expansion (40), where *GID1* repression lines are unable to expand as much as control lines. Taken together, these results demonstrate that SynTF-based repression of *GID1* reduces organ size consistently and in an expression-dependent manner across a range of tissues and developmental times.

### Reductions in organ size generated by SynTF-based repression of *GID1* are stable across fluctuating environments

Thus far, all reported phenotyping had been conducted in controlled growth conditions to minimize potential confounding impacts of variable environments on GA signaling (27). However, as our eventual goal is to use these tools to design crops that are grown in field conditions, it was important to test if the dwarfing generated by SynTF-based repression of *GID1* was stable in fluctuating environmental conditions. For these experiments, three *GID1* repression lines that showed a range of dwarfing strengths in prior experiments and the no gRNA control population (n=5) were grown in tandem in a greenhouse without any active temperature or light control. We chose to quantify dwarfing based on measurements of inflorescence height due to both the ease of measurement of this phenotype in greenhouse grown plants as well as previous reports that *GID1* knockouts show reductions in inflorescence height (17).

Multiple inflorescence phenotypes were observed in the *GID1* repression lines when compared to the no gRNA control lines. Upon phenotyping, we noted a significant delay in the bolting of *GID1* repression lines compared to the no gRNA control lines (**Fig. 2J**), which is consistent with an increased DELLA protein concentration. This provides secondary evidence that the modulation of *GID1* expression is leading to the predicted changes in DELLA protein levels. To prevent differential bolting times from confounding the assessment of inflorescence height, heights were measured once all plants had bolted and were normalized by the number of days post bolting. We observed a significant reduction in inflorescence height in *GID1* repression lines compared to no gRNA control lines (**Fig. 2K**), as predicted. Additionally, the relative strength of dwarfing observed across these lines was consistent with what was observed across experiments in more controlled conditions as inflorescence heights were strongly correlated with both hypocotyl length (r = 0.74, *p* < 0.001, **Fig. S3A**) and root length (r = 0.74, *p* < 0.001, **Fig. S3B**). These results show that SynTF-based repression of *GID1* can generate dwarfing phenotypes that are both consistent across a range of tissues and resilient to fluctuating environments.

### Increasing gRNA recruitment motifs does not increase SynTF repression strength

Our results thus far demonstrate that titrating the strength of *GID1* repression is an avenue to modulate the strength of dwarfing achieved. However, this titration was a product of the inherent variation in T-DNA insertion and copy number associated with transgenesis. We next set out to explore if expression could be more predictably tuned via modifications to the SynTF, specifically via addition of repression domain recruitment motifs at the 3’ end of the gRNA, which has been shown to increase SynTF regulation strength in the context of activators (36). We generated *A. thaliana* transgenic lines containing the same SynTF components but with gRNAs that contain two recruitment motifs (2xPP7) for the repressor (**Fig. S4A**). We then measured *GID1* expression levels as well as organ size, specifically in hypocotyl and rosette tissue, across these lines in tandem with the original lines containing a single recruitment motif gRNA (1xPP7) and no gRNA control lines described above. While we do see significant repression of *GID1* across these lines, we did not see a consistent increase in repression of *GID1* in the 2xPP7 lines compared to the 1xPP7 lines (**Fig. S4B**). Similarly, while we did observe significant dwarfing in these lines, the range of organ size changes across lines was similar to what we observed for the 1xPP7 lines (**Fig. S3C, D**). We hypothesize this may be due to the mode repression implemented by the repressor, TPL, not being amenable to stacking or because the difference in strength of repression achievable via this kind of change to SynTF structure is smaller than the differences that arise from random integration of T-DNAs. These results demonstrate that screening across lines may be a better approach than modifying SynTF structure for tuning regulation strength.

### Activating *DELLA* expression generates similar phenotypes to repressing *GID1*

We next used our model of GA-signaling in roots to interrogate how SynTF-based activation of *DELLA* expression would alter DELLA protein levels in the context of its negative transcriptional feedback. Simulations predicted that the activation of *DELLA* expression would result in higher DELLA protein concentrations and hence a reduction in organ size, i.e., root length, and that the strength of dwarfing would depend on regulation strength (**Fig. S5A**). To test this prediction, we built transgenic *A. thaliana* lines expressing the same SynTF components as used in the *GID1* repression lines as well as gRNAs that were designed to assemble an activator, DREB2A, at the promoters of each of the five *DELLA* genes in *A. thaliana*, namely REPRESSOR OF GA (*RGA*, AT2G01570), GIBBERELLIC ACID INSENSITIVE (*GAI*, AT1G14920), RGA-LIKE 1 (*RGL1*, AT1G66350), RGA-LIKE 2 (*RGL2*, AT3G03450), and RGA-LIKE 3 (*RGL3*, AT5G17490) (**Fig. S5B**).

The primary root lengths of eight-day-old transgenic lines were measured and compared to the same population of no gRNA control lines as described earlier. Of the six independent lines tested, five showed a significant reduction in primary root length compared to controls, spanning a 0.20-to-0.56-fold change, which is consistent with model predictions (**Fig. S5C**). We also observed a significantly higher expression of the two major *DELLAs* targeted by the SynTF, *RGA* and *GAI*, in a line with stronger dwarfing compared to a line with weaker dwarfing (**Fig. S6A, B**). This validates the model’s prediction that the degree of dwarfing is dependent on SynTF regulation strength. Additionally, when we compared the fold change in root length of all the *DELLA* activation lines to that of all the *GID1* repression lines we observed no significant difference in the overall dwarfing generated by these two regulatory architectures (**Fig. S6C**). Taken together, these results demonstrate that activating *DELLA* is a viable alternative to repressing *GID1* for controlling root organ size.

### SynTF-based modulation of *GID1* or *DELLA* expression enables increasing organ size

In the experiments described thus far, we have utilized model-guided reprogramming of GA-signaling gene expression to decrease organ size. However, in some contexts it might be useful to increase organ size, for example increasing fruit size to improve yields or increasing root network size to enable drought resilience. We next explored whether modulating the expression of *GID1* and *DELLA* in the opposite way, i.e., activation of *GID1* expression and repression of *DELLA* expression, can generate organ size increases. We used a modified version of the model described above, with terms for either SynTF-based activation of *GID1* expression or repression of *DELLA* expression, to simulate the DELLA protein concentration at steady state for a range of different SynTF regulation strengths. The model predicted both regulatory architectures would lead to lower concentrations of DELLA protein, and hence an increase in organ size, compared to the no gRNA control (**Fig. 3B, C**). Once again, the magnitude of organ size increase was predicted to be proportional to the strength of regulation.

**Figure 3.**
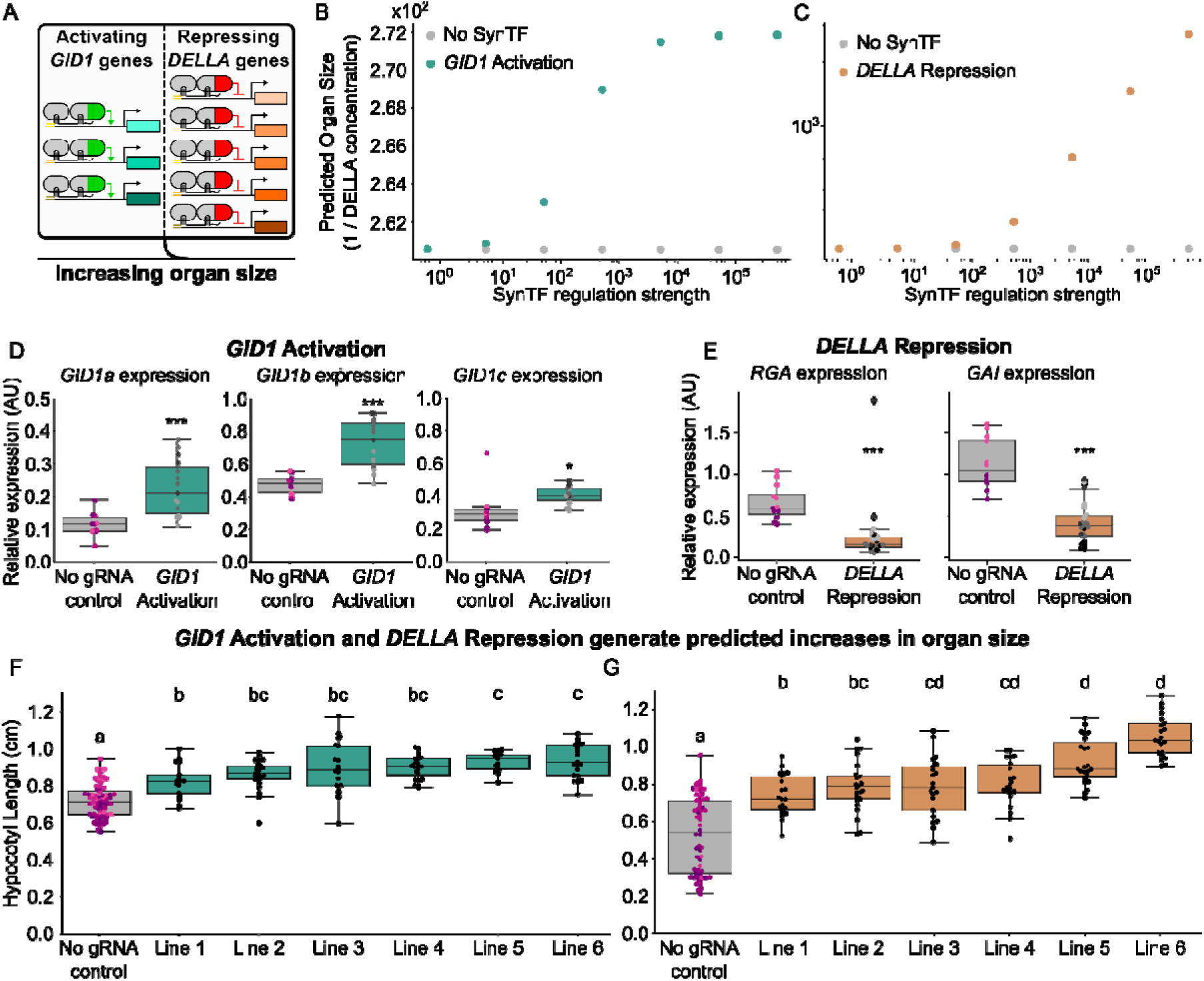
Modeling predicts SynTF regulatory architectures that result in an increase in organ size. A) Schematic describing genetic circuits designed to either activate *GID1* expression or repress *DELLA* expression to increase organ size. B) Plot summarizing the predicted reciprocal of DELLA protein concentration at steady state, used as a proxy for organ size, from simulations of the GA signaling network with either no SynTF (gray) or an activator targeted to regulate *GID1* expression (dark teal), across a range of different SynTF regulation strengths. C) Plot summarizing the predicted reciprocal of DELLA protein concentration at steady state, used as a proxy for organ size, from simulations of the GA signaling network with either no SynTF (gray) or a repressor targeted to regulate *DELLA* expression (dark orange) across a range of different SynTF regulation strengths. D,E) Boxplots summarizing the expression of *GID1a*, *GID1b*, and *GID1c* (D) or *RGA* and *GAI* (E) normalized by housekeeping gene expression (*PP2A*) in seedlings of a population of no gRNA control lines (gray, n=3 and 2, respectively) and either *GID1* activation lines (D, dark teal, n=3) or *DELLA* repression lines (E, dark orange, n=3). F) Boxplots summarizing the hypocotyl length of plant lines with SynTF-based activation of *GID1* expression (dark teal) compared to a population of no gRNA control plant lines (n=4, gray). G) Boxplots summarizing the hypocotyl length of plant lines with SynTF-based repression of *DELLA* expression (dark orange) compared to a population of no gRNA control plant lines (n=4, gray). Every dot of the same color within each boxplot corresponds to an independent biological replicate of the same genotype. Different letters represent statistically significant differences (One-way ANOVA followed by Tukey HSD test, *p* < 0.05). Asterisks represent results of a Welch’s two sample *t*-test (*p* < 0.05), * corresponds to *p* < 0.05, ** corresponds to *p* < 0.005, and *** corresponds to *p* < 0.0005.

To test these predictions, we generated transgenic *A. thaliana* lines expressing the same SynTF components discussed above as well as gRNAs targeting either the three *GID1s* for activation or the five *DELLA*s for repression (**Fig. 3A**). Six independent lines of each genotype were grown in tandem with no gRNA control lines on MS agar plates and hypocotyl length was measured as described above (**Fig. 3F, G**). We also collected RNA from a subset of these lines and measured expression of the target genes. Specifically, we measured the three *GID1s* in the *GID1* activation lines and the two major DELLAs, *RGA* and *GAI*, in the *DELLA* repression lines. We observed significant activation of the three *GID1* genes (**Fig. 3D**), as well as significant repression of *RGA* and *GAI* (**Fig. 3E**) compared to the no gRNA controls across multiple lines. We also observed that all lines tested from both regulatory architectures resulted in a significant increase in hypocotyl length compared to the no gRNA controls with an array of phenotype strengths, ranging from 1.14 to 1.30 and 1.39 to 1.93 fold change over the no gRNA controls for *GID1* activation and *DELLA* repression, respectively, as predicted by the model (**Fig. 3F, G**). We observed a correlation between the total *GID1* or *DELLA* expression and the organ size measured (*GID1* activation: r = 0.52, *p* < 0.001, *DELLA* repression: r = –0.51, *p =* 0.007), validating the predicted relationship between regulation strength and organ size (**Fig. S7A, B**). These results show that model-guided tuning of expression of genes in the GA signaling pathway using SynTFs is a viable approach to engineer both organ size increases as well as decreases.

### Modeling can identify optimal regulatory architectures to engineer organ size

Our results demonstrate that there are multiple ways to modulate the GA signaling pathway to generate a desired organ size change. While there is variation in the relative organ size change, we observe across lines based on the strength of regulation implemented, it is possible that the gene family targeted dictates maximal possible change based on the structure of the GA signaling pathway. However, the feedback loops in the pathway (22) make it challenging to predict the relative impacts of different regulatory architectures, for example *DELLA* repression vs *GID1* activation, on organ size to identify the optimal engineering strategy. We hypothesized that the that the models described above could be leveraged to overcome this challenge by enabling *in silico* prediction of how different SynTF-based regulatory architectures would impact final DELLA protein concentration.

We used our model to interrogate the relative affects that *DELLA* repression and *GID1* activation would have on increases in organ size across different SynTF regulation strengths. Interestingly, the model predicted that *GID1* activation would result in both a smaller linear range in organ size across SynTF regulation strengths, with an earlier saturation, as well as a lower magnitude of organ size increase than *DELLA* repression (**Fig. 4A, S7C**). This prediction is evident when comparing the average hypocotyl length across the six lines of each genotype (**Fig. 3F, G**). We observe a significantly smaller range of hypocotyl length across the *GID1* activation lines (∼0.1 cm) compared to *DELLA* repression lines (∼0.32 cm). To test the model’s prediction that *DELLA* repression would also generate a larger magnitude of organ size increase than *GID1* activation (**Fig. 4A, S7C**), we compared the fold change in hypocotyl length of the lines with the strongest phenotypes for each genotype. We observed a significantly higher fold change in hypocotyl length compared to the no gRNA control for *DELLA* repression (0.96 median difference in fold change) compared to *GID1* activation (0.29 median difference in fold change, **Fig. 4B**). Thus, while increasing organ size is possible via modulation of both *GID1* and *DELLA* expression, *DELLA* repression can generate a stronger phenotype than *GID1* activation. Taken together these results demonstrate how *in silico* simulation can be used to guide SynTF-based engineering of organ size.

**Figure 4.**
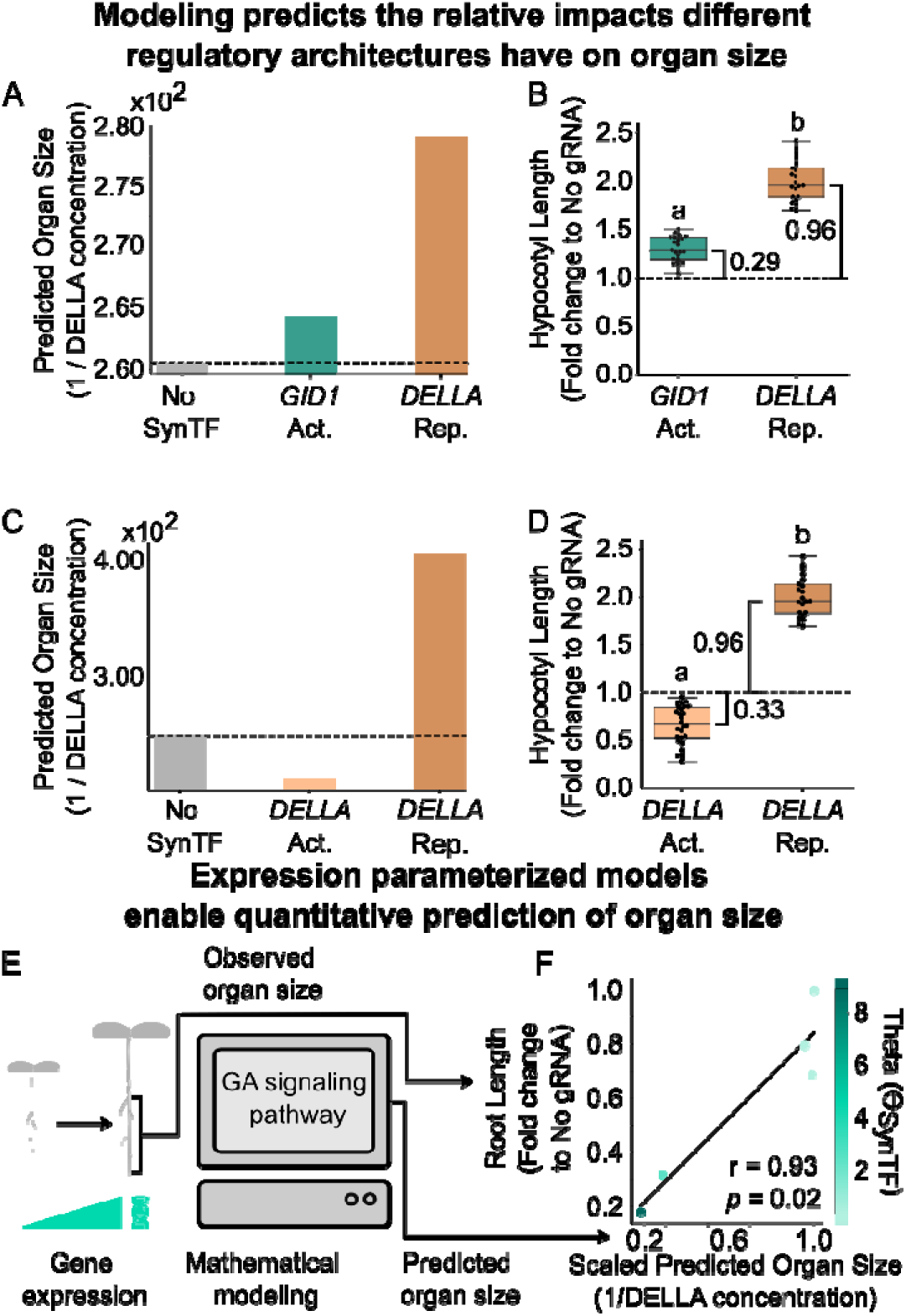
Modeling predicts the relative impacts different regulatory architectures have on organ size. A) Bar plots summarizing the predicted reciprocal of DELLA protein concentration at steady state, used as a proxy for organ size, from simulations of the GA signaling network with either no SynTF (gray), an activator targeted to regulate *GID1* expression (dark teal), or a repressor targeted to regulate *DELLA* expression (dark orange). B) Boxplots summarizing the fold change in hypocotyl length compared to the no gRNA controls of the *GID1* activation (dark teal) or *DELLA* repression (dark orange) line with the strongest phenotype. The brackets represent the difference in fold change of hypocotyl length compared to the no gRNA controls. C) Bar plots summarizing the predicted reciprocal of DELLA protein concentration at steady state, used as a proxy for organ size, from simulations of the GA signaling network with either no SynTF (gray), an activator (orange), or a repressor (dark orange) targeted to regulate *DELLA* expression. D) Boxplots summarizing the fold change in hypocotyl length compared to the no gRNA controls of the *DELLA* activation (orange) or *DELLA* repression (dark orange) line with the strongest phenotype. The brackets represent the difference in fold change of hypocotyl length compared to the no gRNA controls. E) Schematic describing the process of developing expression parameterized models via utilizing gene expression measurements of target genes (*GID1s*) to predict organ size. F) Scatterplot summarizing the Pearson’s correlation between scaled predicted organ (1/DELLA concentration) size generated from ⍰_SynTF_ values of the *GID1* repression lines to the observed mean primary root lengths of the *GID1* repression lines. The color of each dot corresponds to each lines ⍰_SynTF_ value as depicted by the heat map to the right of the plot. The brackets represent the difference in fold change of hypocotyl length compared to the no gRNA controls. In all plots, the dashed black line highlights the phenotype of the no gRNA control. Letters represent statistically significant differences (One-way ANOVA followed by Tukey HSD test, *p* < 0.05)

The further test the predictive power of our model, we used it to investigate whether increasing or decreasing organ size would lead to larger magnitudes of changes. Our simulations predicted that modulating *GID1* or *DELLA* expression to increase organ size would lead to larger magnitudes of change compared to the no gRNA control phenotype than changes in expression that decreased organ size. Specifically, the model predicted that for a given strength of SynTF regulation, targeting *DELLA*s for repression and *GID1s* for activation would generate a larger magnitude in organ size change compared to *DELLA* activation and *GID1* repression, respectively (**Fig. 4C, S8A**). To test this prediction, we first compared the fold change in hypocotyl length of the *DELLA* activation and *DELLA* repression lines compared to the no gRNA controls in the line with the strongest phenotype for both genotypes. We observed that the relative change in magnitude of the *DELLA* repression lines was larger (0.96 median difference in fold change) than the *DELLA* activation lines (0.33 median difference in fold change, **Fig. 4D**), which validates the model’s prediction. Similarly, we tested the model prediction that *GID1* repression would generate a larger change in organ size than *GID1* activation (**Fig. S8A**). We observed a larger fold change in hypocotyl length compared to the no gRNA controls when activating *GID1*s (0.29 median difference in fold change) than when repressing *GID1*s (0.26 median difference in fold change) (**Fig. S8B**). These results demonstrate the capacity of *in silico* modeling to predict the relative impact modulating different nodes in the GA signaling pathway.

### Expression parameterized models enable quantitative prediction of organ size

In all of the models generated in this work so far, we have assumed a SynTF regulation strength that approximately matches the regulation strength of the DELLA, and thus they are limited to qualitative predictions of the relative impact of different regulatory architectures. However, our empirical measurements of the strength of regulation on SynTF-targeted genes across different lines provide an avenue to move towards more quantitative predictions of impacts on organ size (**Fig. 4E**). Specifically, we assume that as these lines are near isogenic aside from a different SynTF regulation strength (denoted as ⍰_SynTF_), we hypothesize that we could fit this parameter based on our measurements for each line to create an individualized model capable of quantitative prediction of organ size changes relative to the no gRNA controls.

To test this, we focused on four lines that span a range of SynTF regulation strength in our best characterized genotype, the *GID1* repression lines. Here, we used the fold change of the sum of *GID1a* and *GID1c* expression compared to the mean of the no gRNA control population to fit a ⍰_SynTF_ for each line (**Table S1**). *GID1b* was not considered as we do not see significant repression of this gene. Subsequently, we simulated the steady state DELLA protein concentration for each line’s ⍰_SynTF_ and used its reciprocal to determine the predicted fold change in organ size relative to the no gRNA control. A more detailed description of this calculation and the code used to implement it can be found in the methods section. These predicted sizes were then compared to the observed measurements of mean primary root length of each of these lines, as this is the tissue that the rest of model parameters were originally based on, and were found to be strongly correlated (r = 0.93, p = 0.02, **Fig. 4F**). This validates the utility of the expression parameterized approach to enable quantitative predictions of changes in organ size.

We next explored if these expression parameterized models could facilitate prediction of how the organ size changes generated by lines with different SynTF regulation strengths would respond to environmental fluctuations that might occur in an agricultural context. It has been shown that fluctuations in temperature result in changes in endogenous GA levels via modulation of GA biosynthesis genes (41). To explore how this would impact the SynTF-mediated changes on organ size, we built modified versions of the models described above where GA concentration was fixed at a low or high level and then simulated the steady state DELLA protein concentration to predict organ size, as above. These simulations were performed for three *GID1* repression lines (**Fig. S9A**) as well as three *GID1* activation lines (**Fig. S9D**) whose ⍰_SynTF_ values were calculated as described above.

The models predicted an overall increase in organ size across all lines at the higher temperature associated with the higher GA concentrations leading to lower DELLA protein concentrations. Interestingly, our models also predicted that the fold change in organ size relative to the no gRNA control would behave differently for *GID1* repression lines (**Fig. S9A, B**) versus *GID1* activation lines (**Fig. S9D, E**). Namely, the fold change in organ size of *GID1* repression lines relative to the no gRNA control would decrease (more similar in size to the no gRNA controls) at higher temperatures (27⁰C), with lines that have a higher ⍰_SynTF_ value having a smaller reduction in size (**Fig. S9B**). In contrast, the fold increase in organ size of *GID1* activation lines relative to the no gRNA control is almost unchanged at high temperatures across all ⍰_SynTF_ values (**Fig. S9E**).

To test these predictions, we grew three *GID1* repression, three *GID1* activation, and three no gRNA control lines on MS agar plates in both low (22⁰C) and high (27⁰C) temperatures to simulate low and high GA environments. We then measured hypocotyl length across all lines. At high temperature, hypocotyls significantly increased in size across all lines (**Fig. S10**), as predicted. The overall temperature-induced alteration to fold change in hypocotyl length was also larger in the *GID1* repression lines (0.17 mean fold change in lines that showed *GID1* repression) than the *GID1* activation lines (0.05 mean fold change; **Fig. S9C, F**). Specifically, *GID1* repression lines with a larger ⍰_SynTF_ resulted in larger fold change compared to the no gRNA controls at higher temperatures. For example, the three *GID1* repression lines with ⍰_SynTF_ values of 0.01, 0.44, and 9.28 corresponded to a fold change of 0.02, 0.08, and 0.26, respectively. Finally, the temperature-induced alteration to fold change in hypocotyl length in the *GID1* activation lines was minimal. The *GID1* activation lines with ⍰_SynTF_ values of 2.12, 38.3, and 30.7 corresponded to a fold change of 0.09, 0.04, and 0.03, respectively, as predicted by the model. We simultaneously performed an assay with the same lines on MS agar plates with low (10 µM) and high (100 µM) GA_3_ to validate that the effect we observed is from changes in GA levels, and observed similar results and trends as observed in the temperature assay (**Fig. S11**). Taken together, these results demonstrate the utility of expression parameterized models to quantitatively predict the strength and stability of organ size phenotypes.

### SynTF-based repression of *GID1s* can generate expected phenotypes in tomato

Our results thus far demonstrate that model-guided SynTF-based expression reprogramming of GA-signaling genes enables predictable modulation of organ size in the model species, *A. thaliana*. However, as the goal of this work is to elucidate the design rules for engineering organ size in crops, we next explored whether these principles could be generalized to a major crop plant, *Solanum lycopersicum* (tomato). Specifically, we focused on testing whether SynTF-based repression of *GID1* could generate robust reduction in organ size across tissues in tomato as such compact morphologies have the potential to increase yields in indoor growth environments where space is at a premium (42).

We generated transgenic tomato lines with a single protein SynTF, consisting of a nuclear localized deactivated Cas9 fused to the N-terminal 188 amino acids of the TOPLESS repressor (34), and gRNAs targeting the promoters of the three *GID1* genes in tomato, *GID1a* (Solyc01g098390), *GID1b-1* (Solyc09g074270), and *GID1b-2* (Solyc06g008870, **Fig. 5A**). These lines were grown in tandem with controls lacking the SynTF and the length of hypocotyls was measured. We observed a significant decrease in hypocotyl length (0.61-to-0.84-fold change) across all five independent *GID1* repression lines tested when compared to the no SynTF control lines (**Fig. 5B**). We also measured primary root length across all lines. All *GID1* repression lines showed a decrease in size (0.43-to-0.91-fold change), however, only two of the independent *GID1* repression lines were statistically significant (**Fig. 5C**). This could be due to the small effect size produced in the non-significant lines and small sample sizes. Nevertheless, we still observed that the lines with the strongest shoot dwarfing also had the strongest root dwarfing, replicating the consistent impact across tissues (**Fig. S12**).

**Figure 5.**
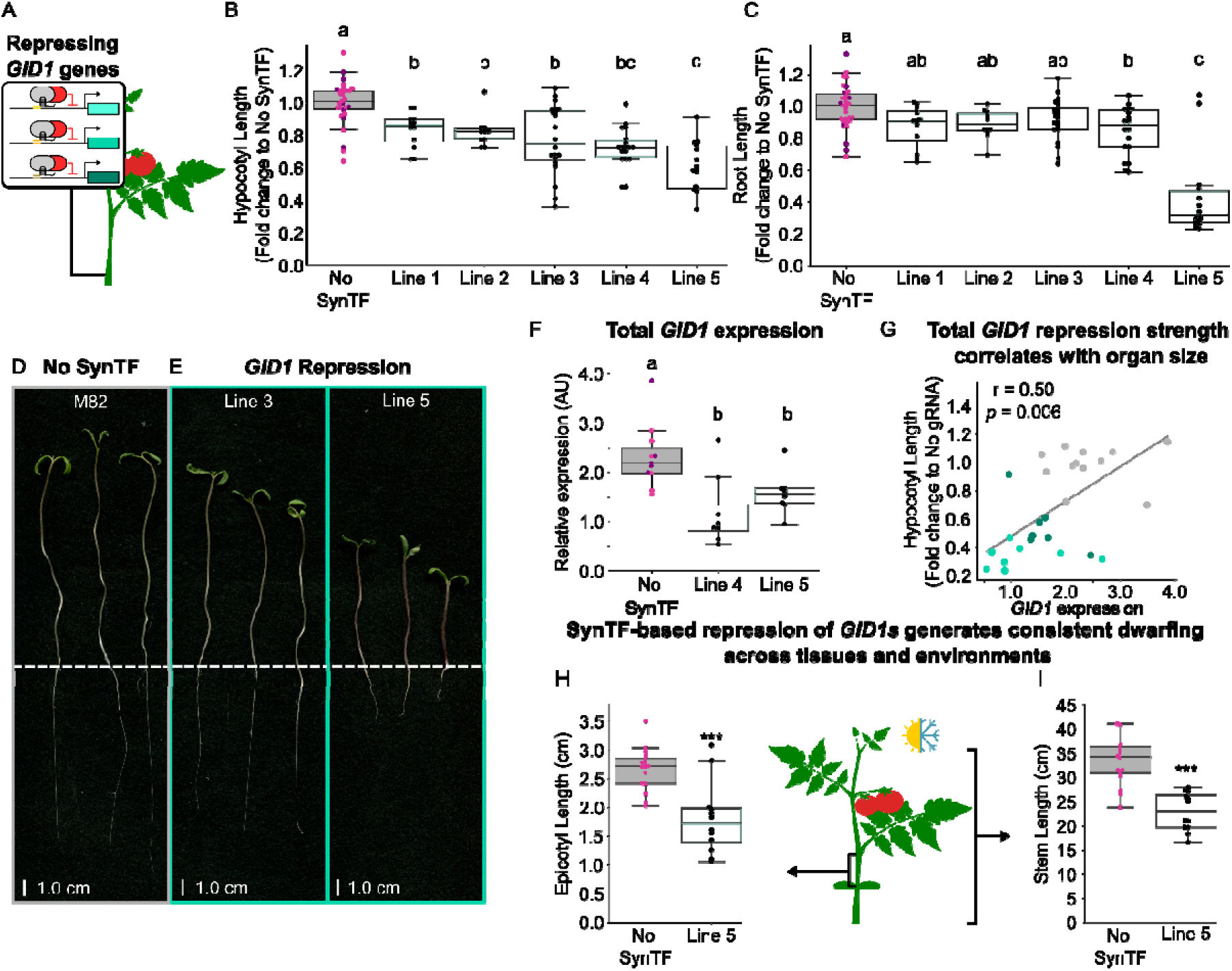
Repression of GID1 generates consistent dwarfing across tissues and fluctuating environments in tomato. A) A schematic of the Cas9-based synthetic transcription factor used to implement repression on the three different *GID1* homologs in *S. lycopersicum* (tomato), which is predicted to decrease organ size. B) Boxplots summarizing hypocotyl length of five independent transgenic lines with SynTF-based repression of *GID1* (teal) compared to control plant lines lacking the SynTF (gray, n=2). C) Boxplots summarizing the fold change in primary root length to the no SynTF lines (gray, n=2) of the five independent *GID1* repression lines (teal). D,E) Representative seedling phenotypes of the no SynTF control (D) and two independent *GID1* repression lines (E). The white dashed lines approximately separate the root and shoot tissue. F) Boxplots summarizing the total *GID1* expression (sum of *GID1ac*, *GID1b*-1, and *GID1b*-2) in the no SynTF lines (gray, n=2) and the two *GID1* repression lines (teal) that showed the strongest hypocotyl phenotype. G) Scatterplot summarizing the Pearson’s correlation (r=0.50, *p* = 0.006) between the fold change in hypocotyl length of the *GID1* repression lines (shades of teal) to the no SynTF lines (gray, n=2) and the total *GID1* expression (sum of *GID1ac*, *GID1b*-1, and *GID1b*-2) H,I) Boxplots summarizing epicotyl (H) and stem (I) length of greenhouse-grown no SynTF controls (gray, n=1) and the *GID1* repression line with the strongest phenotype (teal). Across all plots, every dot of the same color corresponds to an independent biological replicate of the same genotype. Different letters represent statistically significant differences (One-way ANOVA followed by Tukey HSD test, *p* < 0.05). Asterisks represent results from a Welch’s two sample *t*-test (*p* < 0.05), * corresponds to *p* < 0.05, ** corresponds to *p* < 0.005, and *** corresponds to *p* < 0.0005.

To validate the dwarfing phenotypes we observed were due to the repression of *GID1s*, we measured expression of the target genes in hypocotyls of the no SynTF control lines and the two *GID1* repression lines with the strongest hypocotyl phenotype. Both *GID1* repression lines analyzed showed a significant reduction in total normalized *GID1* expression (sum of *GID1ac*, *GID1b*-1, and *GID1b*-2) when compared to the no SynTF controls (**Fig. 5F**). Additionally, we observed that *GID1* repression strength correlates with dwarfing strength in these lines (r = 0.05, *p* = 0.006, **Fig. 5G**), as predicted by the model. This data provides evidence that the model-guided design principles for SynTF-based engineering of organ size can be generalized to tomato.

Finally, we wanted to investigate whether the dwarfing phenotypes we observed were stable in fluctuating environments. We grew the *GID1* repression line with the strongest phenotype and the no SynTF control in a greenhouse with no active light or temperature control and quantified both epicotyl (**Fig. 5H**) and stem length (**Fig. 5I**). The *GID1* repression line showed consistent dwarfing in both tissues when compared to the no SynTF control. This, once again, demonstrates that SynTF-based repression of *GID1* results in consistent dwarfing across tissues and fluctuating environments and provides evidence that insights we generated in *A. thaliana* can be generalized across species.

## Discussion

Our results demonstrate that repression of the *GID1* genes with SynTFs can generate predictable and tunable dwarfing in *A. thaliana* roots. We observed that *GID1* repression generated a range of reductions in root lengths across independent lines that correlated with the strength of SynTF-implemented repression of *GID1*. This was consistent with the model’s prediction that increasing strength of repression by the SynTF would result in higher DELLA protein concentrations and hence larger reductions in organ size. While prior studies had used knockouts and overexpression to elucidate the role that *GID1* genes play in regulating cell expansion (13–15), our results are the first demonstration that titrating their expression is sufficient to generate predictable, dosage-dependent changes in organ size (**Fig. 1E-G**). One limitation of our study is the non-functionality of the *GID1b* gRNA, resulting in incomplete control of total *GID1* expression. Future studies using viral-based gRNA screens (11) could enable identification of gRNAs that facilitate co-regulation of all three *GID1* genes, potentially increasing the strength of dwarfing generated.

The approach of tuning the expression of this gene family to generate changes in organ size will enable future interrogation of the relationship between organ size and physiology beyond the extremes generated by knockouts and overexpression, and without confounding co-variation in other traits that can stymie studies of natural variation. Our observation that the repression implemented by SynTFs was only visible in tissues collected in the evening is consistent with the reported circadian oscillations of *GID1* expression (37). This suggests a molecular model where, in the no gRNA controls, the spike of *GID1* expression in the evening reduces DELLA protein levels and enables cell expansion, while in the *GID1* repression lines this spike is suppressed leading to lower cell expansion.

This work also shows that *GID1* repression enables reduction of organ size across tissues and environments. This demonstrates that the GA-signaling based control of cell expansion functions similarly across cell types and implies that SynTF-based expression reprogramming can be applied to engineer the size of a range of tissues. We also observed significant dwarfing in *GID1* repression lines grown in variable greenhouse environments, which indicates that these circuits are stable in fluctuating environmental conditions. This is a critical determinant of the utility of this approach for engineering crops, which are grown in variable field and greenhouse environments.

The comparison between the reduction in root length observed in *A. thaliana* lines with *DELLA* activation and *GID1* repression demonstrates that *DELLA* activation is a viable alternative for generating dwarfing. This is a valuable insight as the *GID1* and *DELLA* gene families have different numbers of homologs across species, for example *A. thaliana* has five DELLAs while tomato has one. Because SynTF regulation tends to become less efficient with more targets due to competition for resources, targeting the minimum number of genes is desirable for implementing strong regulation. Thus, knowing that regulation of either *DELLAs* or *GID1s* can generate equivalent dwarfing enables selecting the smallest gene family to target in a crop.

In addition to reducing organ size, our results also demonstrate that modulating expression of *GID1* and *DELLA* in the opposite direction is an avenue to increase organ size. While both *GID1* activation and *DELLA* repression generated increases in *A. thaliana* hypocotyl length, we observed a significantly larger effect on organ size from *DELLA* repression overall, which was consistent with model simulations. This may be related to the direct repression of *DELLA* by SynTF being more efficient at decreasing DELLA levels compared to activating *GID1s*, which require an additional partner (bioactive GA) to decrease DELLA levels in the cell. Additionally, DELLAs instigate negative feedback on themselves. Adding a synthetic repressor to the DELLAs may directly modulate their expression and impact the capacity of the native de-repression that occurs at lower DELLA levels, resulting in consistently lower steady state DELLA concentrations. The model also accurately predicted the larger fold change in organ size from *DELLA* repression compared to *DELLA* activation, and from *GID1* activation compared to *GID1* repression, which highlights its predictive power. More broadly, these results demonstrate how such models can be used to compare different SynTF-based regulatory architectures to identify the optimal perturbations for achieving a target morphology, particularly in gene networks containing feedback loops.

We show that parameterizing these models with line specific expression measurements expands their utility from qualitative comparisons at the genotype level to quantitative predictions of phenotype, specifically root length, of individual lines. To our knowledge this is the first demonstration of quantitative prediction of tissue level morphology using models of molecular signaling in plants. The model predicts a sigmoidal relationship between regulation strength and organ size, where there is a linear relationship that becomes static when the regulation strength is above or below a threshold level (**Fig. 1B**). Our expression parameterized models were fit using data from *GID1* repression lines that fell within the linear regime of the model prediction (**Fig. 4F**). Our SynTF architectures were unable to generate regulation strong enough to enter the static regime. In future work we hope to explore alternative SynTF designs to interrogate the, as of yet, unexplored non-linear parts of the regulation strength-organ size relationship. We also show how such models could be applied to identify lines that are best able to maintain consistent phenotypes in the face of environmental fluctuations, potentially enabling consistent performance in variable agricultural environments. Parameterizing similar models for the relevant tissues of crop species will facilitate the precision design of plant morphology using SynTF-based expression reprogramming.

Finally, the demonstration that *GID1* repression can also generate dwarfing across tissues in tomato, implies that the design principles for engineering organ size elucidated above can be generalized across species. This is likely due to the highly conserved role GA-signaling plays in the control of cell expansion across plant species as well as the orthogonal nature of SynTF based regulation (43, 44). However, rigorous validation of this claim will require characterization of the phenotypic outcomes of these circuits across a range of monocot and dicot crop species. These results highlight the promise of utilizing model-guided SynTF-based expression engineering for bottom-up design of morphology across a range of crops in the future.

However, while promising, there are still aspects of this approach that need to be refined to achieve its full potential. While the existing systems are useful for modulating organ size, such global changes are often associated with negative pleiotropies. For example, while dwarfed stems might increase productivity and resilience, associated root dwarfing can reduce water and nitrogen use efficiency. In addition, the cross regulation of other traits by GA signaling, such as flowering time (18, 45, 46), mean global reprogramming of GA signaling could also lead to unintended consequences such as the delayed flowering we observed in our greenhouse experiments. Restricting the activity of SynTFs using tissue specific expression strategies could enable targeted changes to the size of specific organs. This highlights a potential benefit of targeting GA signaling genes, such as *GID1s* and *DELLAs*, over the strategies that have been explored thus far which focus on modulation of GA biosynthesis. As GA is actively trafficked through plant tissues, local changes could result in systemic morphological alterations. In contrast, targeting the GA signaling genes with SynTFs would sidestep these challenges as they have cell autonomous effects, thereby minimizing the chances of transport related pleiotropies.

Additionally, while this work provides a robust synthetic framework for controlling cell expansion, this is only one of several determinants of organ size, which limits the extent of morphological reprograming possible. This was demonstrated in our ability to restrict the upper bound, but not the lower bound of leaf size in *A. thaliana*. Tools for the precision control of cell division and extracellular matrix deposition will be necessary to enable a holistic bottom-up control of organ size in the future. Finally, we show that our approach can generate a range of phenotype strengths, thanks to the range of regulation strengths obtained from insertional variation of the SynTFs across transgenic events. Transitioning from this line generation and screening approach to the predictable tuning of phenotype requires two major innovations. Firstly, strategies to predictably tune the regulation strength of SynTFs and secondly, technologies such as landing pads or targeted insertion into validated safe harbors to enable consistency across transgenic events.

Enabling the bottom-up design of crop traits via precision changes to gene expression would be a step change in the process of crop enhancement. Moving from iterative phenotype-based selection of beneficial natural mutations to model-guided alterations of gene expression using SynTFs would accelerate trait engineering. Additionally, the use of SynTFs that are orthogonal to endogenous signaling pathways enables predictable alterations to gene expression and associated phenotypes across tissues and species, as we demonstrate. This contrasts with *cis*-regulatory variation isolated from breeding approaches that can lead to variable phenotypes across genetic backgrounds thanks to cross-regulation of pathways, epistasis and linkage drag. Finally, bottom-up design with synthetic systems would grant access to regions of biological design space that have not been sampled by natural variation due to evolutionary bottlenecks or associated negative pleiotropies.

Overall, this work advances our fundamental understanding of GA signaling, specifically the relationship between signaling gene expression and organ size. It also lays the groundwork for the bottom-up design of crop morphology by elucidating the design principles for using SynTFs, guided by mathematical models of GA signaling, to reprogram organ size. This opens the door to a range of exciting translational applications including engineering plant-based biomaterials and creating morphologies that maximize productivity in specific environments, be they fields, indoor farms, or spaceships.

## Materials and Methods

### Model simulations

A modified version of the mathematical model of GA signaling, originally developed by Middleton et al. (22), was implemented in python to simulate the qualitative impact of SynTF-based regulation of *GID1* or *DELLA* expression on steady state *DELLA* protein levels. This involved adding equations to simulate SynTF transcription and translation as well as modifications to the Hill equations simulating *GID1* activation or repression or *DELLA* activation or repression (**Fig. S1**, **Table S1**). These equations, their constants, and more information about the mathematical modeling are available in the supplemental materials and methods.

To quantitively predict organ size, we developed expression parameterized models. This involved using the measured *GID1* repression levels to calculate a ⍰_SynTF_ for four *GID1* repression lines, predicting organ size for each line using this ⍰_SynTF_, and finally comparing the measured fold change in root length to this prediction. The expression parameterized models and ⍰_SynTF_ values generated for three *GID1* repression and activation lines were then used to explore how environmentally generated fluctuations of GA levels impact the phenotypes generated in these lines. More information for all of the mathematical modeling is located in the supplemental materials and methods (**Fig. S1**, **Table S1**) and the associated jupyter notebook on github (https://github.com/arjunkhakhar/250610_GA_signaling_paper).

### Plasmid construction

All plasmids constructed in this work were built using Golden Gate Assembly and Modular Cloning (47, 48) and are listed with links to their annotated plasmid maps in **Supplemental Table S2**. All plasmids used in this work will be available via AddGene. More information regarding the DNA parts used and the assembly process is available in the supplemental materials and methods.

### Transgenic line regeneration

The *Arabidopsis thaliana* transgenic lines described in the work were generated by introducing the SynTF components and gRNAs targeting the promoters of either the *GID1s* or *DELLAs* into the genome of Col-0 via floral dip (51). Multiple independent T1 or T2 lines were used for further experiments. The transgenic tomato lines described in this work were generated via construction of a T-DNA encoding the Cas9-based repressor described in the main text as well as gRNAs to target it to the promoters of the three *GID1* genes and two non-functional gRNAs. Subsequent *Agrobacterium*-mediated transformation was outsourced to the plant transformation center at UC Davis. The recovered transgenic lines were confirmed for SynTF expression via RT-qPCR and multiple independent T2 lines were used for subsequent experiments.

### *Arabidopsis* root phenotyping

Seeds were surface sterilized and were plated on 1/2x MS (52) media. Seeds were stratified in the dark at 4⁰C for five days and placed vertically under short day conditions (9hr light/15hr dark) at 22⁰C in a Percival growth chamber for eight days. Plates were scanned using an Epson scanner and root length was measured using ImageJ. More information about the sterilization and seed preparation is available in the supplemental materials and methods.

### Gene expression analysis via RT-qPCR in *Arabidopsis*

For characterization of *GID1* and *DELLA* expression, whole eight-day-old seedlings from the root phenotyping assays of the different regulatory architectures and no gRNA control lines were harvested either during the day or one hour after dark and immediately frozen in liquid nitrogen. RNA was extracted from these tissues and concentrations of *GID1a*, *GID1b*, *GID1c*, *RGA*, *GAI*, and *PP2A* were quantified using RT-qPCR. Cycle threshold values for *GID1* and *DELLA* were normalized to the housekeeping gene (*PP2A*). Two technical replicates were performed for all samples. More information regarding the gene expression analysis pipeline is located in the supplementary materials and methods.

### *Arabidopsis* shoot phenotyping

For hypocotyl measurements, seeds of *A. thaliana* transgenic lines were surface sterilized, plated on media plates, and germinated as described in the supplemental materials and methods. For the experiments investigating GA levels on organ size, GA_3_ was added to the media at concentrations of 10 and 100 µM. Plates were stratified for three days in the dark at 4⁰C, light shocked for six hours in a Percival growth chamber set to 22⁰C or 27⁰C and short-day conditions (9hr light/ 15hr dark) and then were wrapped in foil and placed vertically in the correct growth chamber to promote etiolation. After three days, plates were scanned using an Epson scanner and hypocotyls were measured using Image J.

For rosette and leaf phenotyping, seeds of *A*. *thaliana* transgenic lines were surface sterilized, plated, and germinated as described in the supplemental materials and methods. Twelve, 15-day-old seedlings were transplanted into soil and grown in a Conviron growth chamber as described in the supplemental materials and methods. After 15 days in the growth chamber, each plant was dissected by leaf and scanned in order of developmental age. Individual leaf areas were measured using ImageJ. Rosette areas were calculated by summing the areas of individual leaves for each plant.

For inflorescence phenotyping, seeds of *A. thaliana* transgenic lines were surface sterilized, plated, and germinated as described in the supplemental materials and methods. on 1/2x MS media plates as described above. Three-week-old seedlings were transplanted into soil and grown as described in the supplemental materials and methods. Plants were monitored daily and the date that individual plants transitioned to reproductive growth was recorded. Once all plants had inflorescences that reached at least approximately two centimeters in height, inflorescence height was collected. Inflorescence height was normalized by days-post bolting for better comparisons of dwarfing between genotypes.

### Tomato phenotyping and gene expression analysis

Seeds of M82 WT, M82 Cas9, and *GID1* repression tomato lines were surface sterilized and germinated as described in the supplemental materials and methods. Hypocotyl and root length of 14-day-old seedlings were measured and seedlings were transplanted into ProMix HP soil in 4-inch square pots and placed in a greenhouse with no active temperature or light control. After six days and 34 days, epicotyl and stem lengths were measured, respectively.

To measure *GID1* expression in tomato plants, hypocotyl tissue was collected one hour after dark by sampling 14-day-old seedlings. The concentrations of *GID1a*, *GID1b-1*, *GID1b-2,* and *CAC* were quantified using RT-qPCR. Two technical replicates were performed for all samples. Cycle threshold values of the *GID1* genes were normalized to the housekeeping gene (*CAC*). More information regarding the gene expression analysis pipeline is available in the supplemental materials and methods.

### Data analysis and plotting

All data analysis was performed in python using Jupyter Lab (version 3.4.4). The *p*-values reported were calculated using either an ANOVA followed by a Tukey HSD test or Welch’s *t*-test for pairwise comparisons, where appropriate using the scipy package in python. All the data that is presented was plotted using the seaborn package and functions from the matplotlib library (54). All the raw data and code used to analyze the data is available on the following github repository. (https://github.com/arjunkhakhar/250610_GA_signaling_paper).

## Acknowledgments

We would like to thank Dan Voytas and Colby Starker for their help with transgenic lines and plasmids. We would also like to thank Sophia Seib and Justin Bull for assistance in phenotyping. T.B. and J.V.B. were supported by an award from the CROPPS center funded by National Science Foundation Grant No. DBI-2019674. A.K. was supported in part by funding from the National Science Foundation under Grant No. 2310396.

## Author Contributions

T.B. and A.K. designed the research. T.B., J.V.B., S.S., and H.C. performed the experiments. T.B. and A.K. wrote the manuscript.

## Classification

Plant synthetic biology.

## Supplementary Information

### Supplementary Figures

**Supplementary Figure S1.**
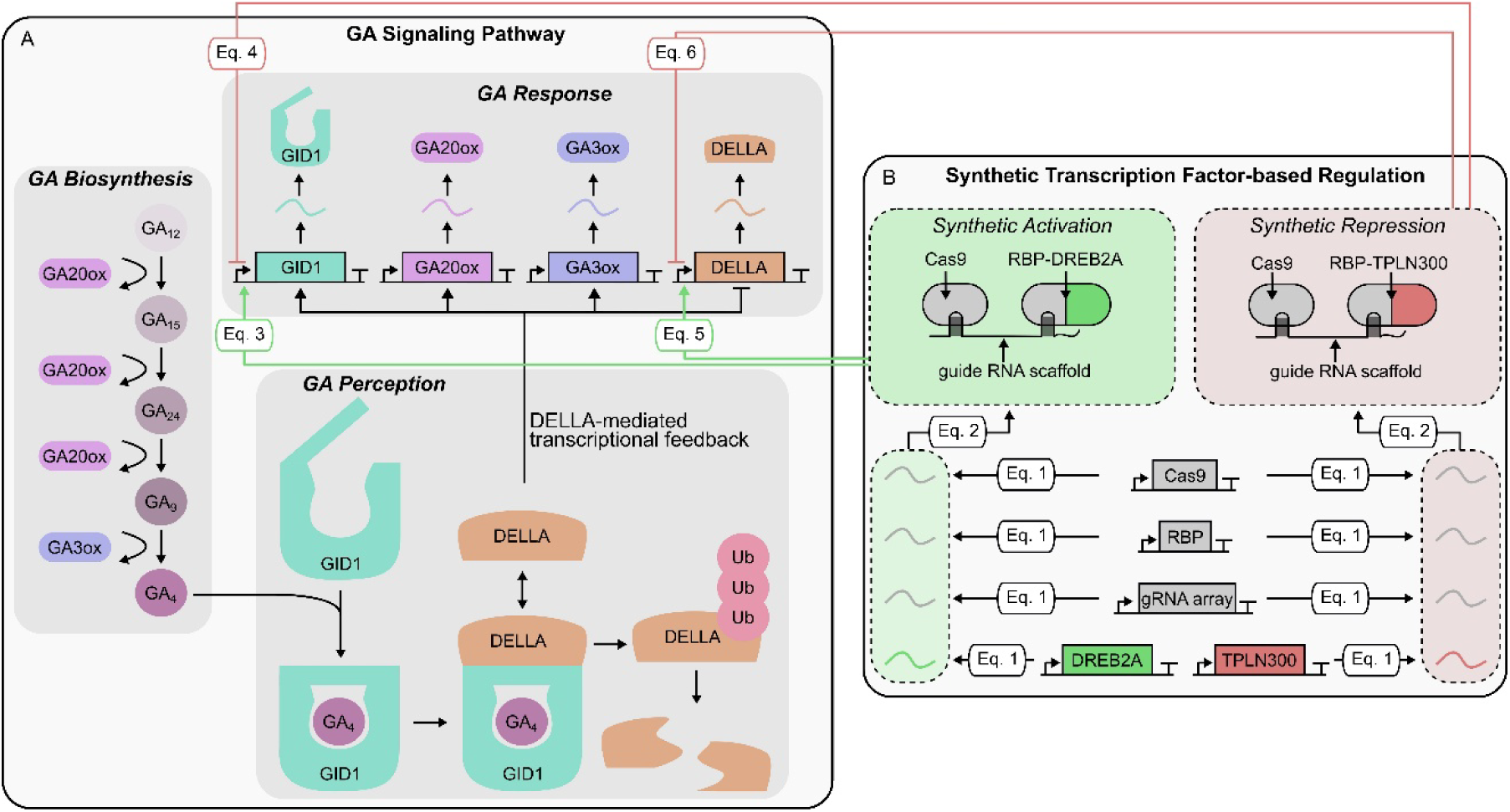
The GA signaling pathway and implemented synthetic transcription factor-based regulation. A) Schematic of the GA signaling pathway depicting all modeled components and the DELLA-mediated feedback signaling native to the pathway. B) Schematic depicting the implemented SynTF-based proteins for targeted activation or repression of *GID1s* and *DELLAs*. The white boxes represent the ODE equations that simulate transcriptional regulation of *GID1s* and *DELLAs* with the SynTF. The equations are listed in the supplementary methods.

**Supplementary Figure S2.**
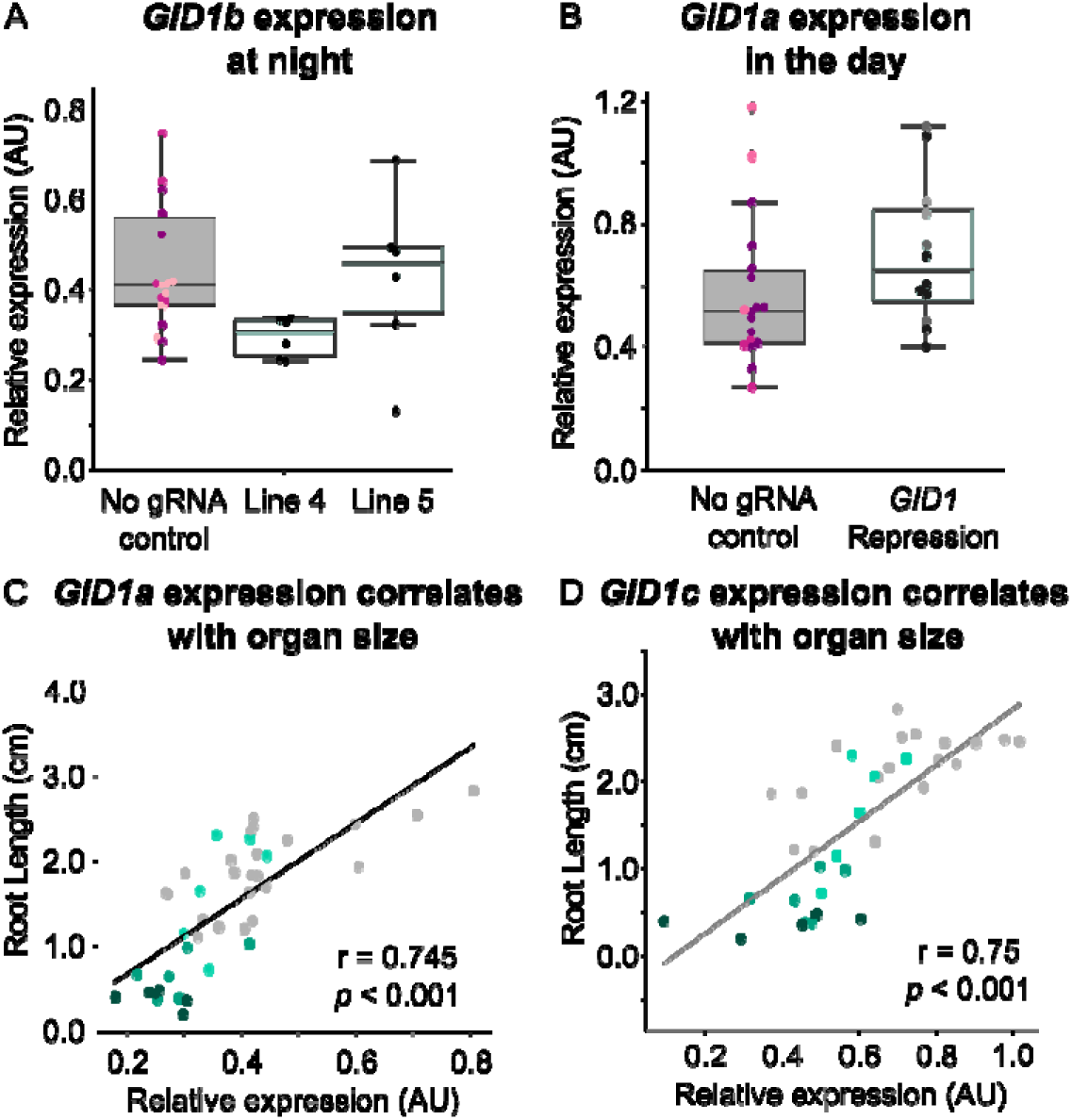
A) Boxplots summarizing the expression of *GID1b* normalized by housekeeping gene expression (*PP2A*) in seedlings of a population of no gRNA control lines (gray, n=3) and the *GID1* repression lines with the strongest phenotypes (teal). Every dot of the same color in each boxplot corresponds to an independent biological replicate of the same genotype. B) Boxplots summarizing the expression of *GID1a* normalized by housekeeping gene expression (*PP2A*) in seedlings whose tissue was collected in the day rather than in the evening as in Fig. 1, from a population of no gRNA control lines (gray, n=3) and the *GID1* repression lines with the strongest phenotypes (teal). Every dot of the same color corresponds to an independent biological replicate of the same genotype. C,D) Scatterplots that depict the Pearson’s correlation between root length and normalized *GID1a* (C) and *GID1c* (D) expression. Every dot of the same color corresponds to an independent biological replicate of either the no gRNA control lines (gray) or *GID1* repression lines 1, 4, and 5 (progressively darker shades of teal).

**Supplementary Figure S3.**
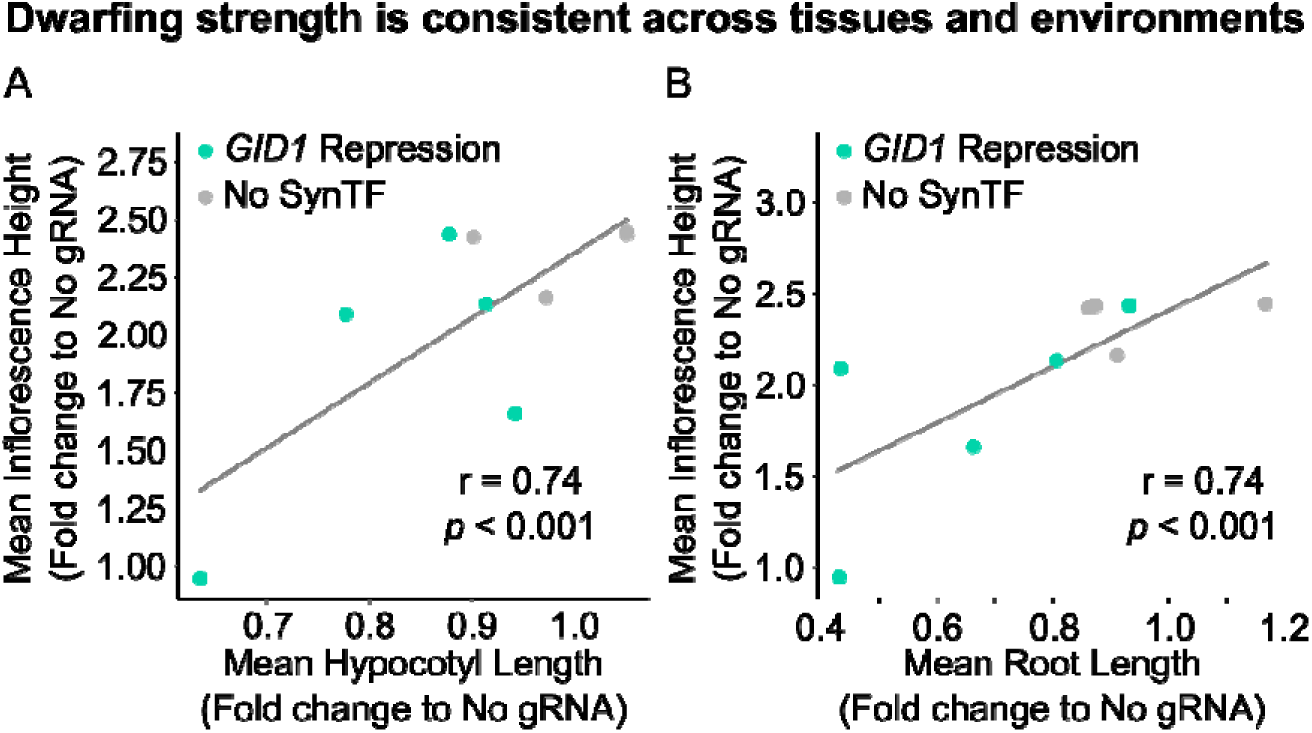
Dwarfing is consistent across tissues and environments. A,B) Scatterplots that depict Pearson’s correlation between mean fold change in inflorescence height and either mean fold change in hypocotyl length (A) or mean root primary root length (B) of *GID1* repression lines (teal) and no gRNA control lines (gray). Fold change is quantified as fold change to the no gRNA controls.

**Supplementary Figure S4.**
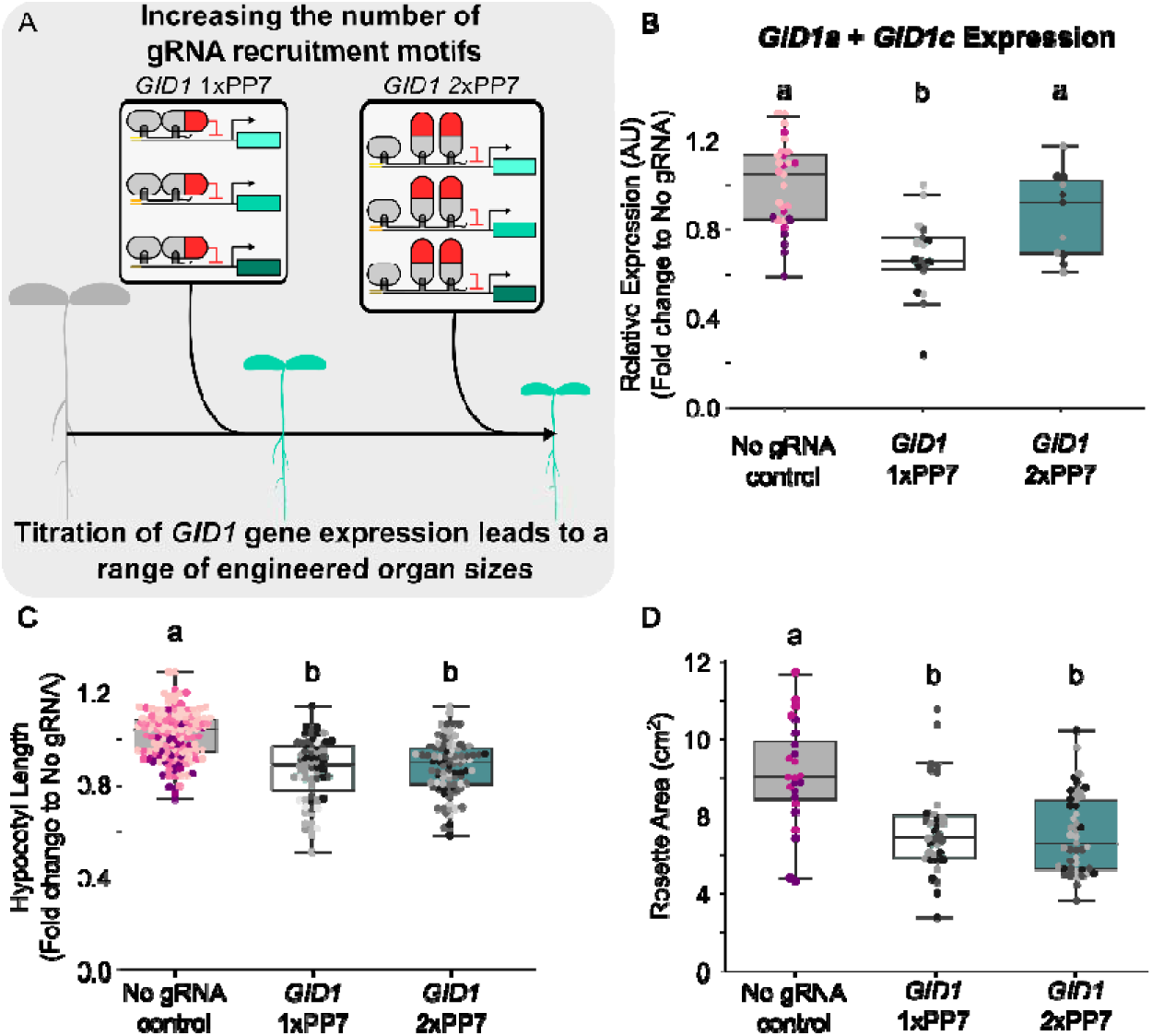
Attempted increase of SynTF regulation strength via incorporation of multiple gRNA recruitment motifs. A) Graphic depicting the recruitment motif engineering strategy and hypothesized organ size changes. B) Boxplots summarizing the *GID1* expression levels (sum of *GID1a* and *GID1c* expression) of the no gRNA controls (gray, n=4), the *GID1* 1xPP7 repression population (light teal, n=3), and the *GID1* 2xPP7 repression population (dark teal, n=2). C,D) Boxplots summarizing hypocotyl (C) and rosette area (D) phenotypes for the no gRNA control population (gray, n=7), the *GID1* 1xPP7 repression population (light teal, n=5), and the GID1 2xPP7 repression population (dark teal, n=6). Every dot of the same color within each boxplot corresponds to an independent biological replicate of the same genotype. Different letters represent statistically significant differences (One-way ANOVA followed by Tukey HSD test, *p* < 0.05).

**Supplementary Figure S5.**
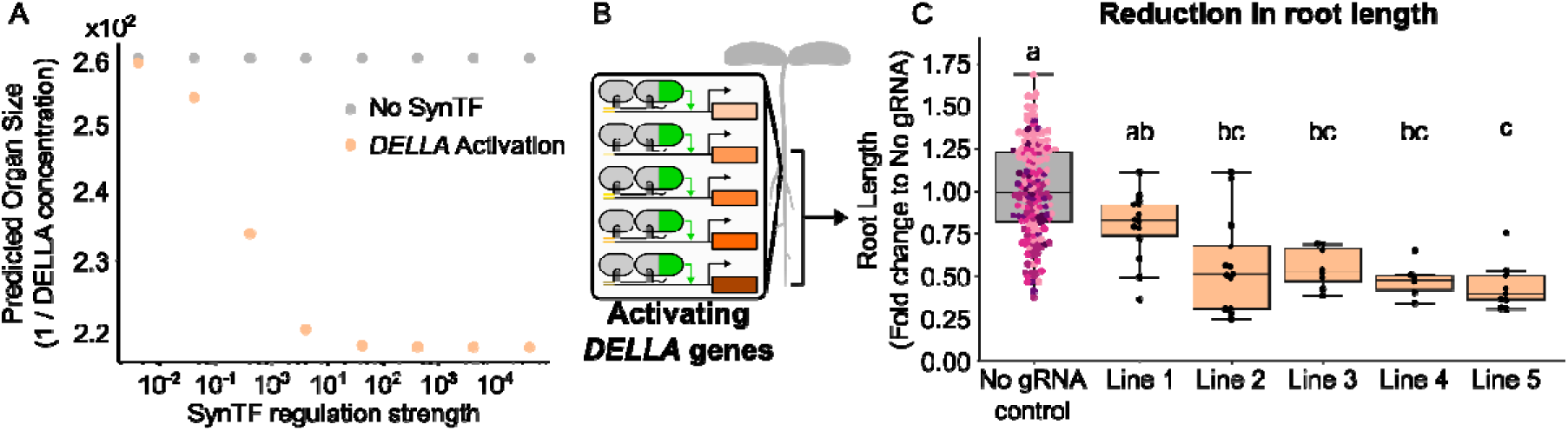
SynTF-based activation of DELLA genes generates model-predicted dwarfing. A) Plot summarizing the predicted reciprocal of DELLA protein concentration at steady state, used as a proxy for organ size, from simulations of the GA signaling network with either no SynTF (gray) or an activator targeted to regulate *DELLA* genes (orange), across a range of different SynTF regulation strengths. B) Schematic of the genetic circuit designed to implement SynTF-based activation on five *DELLA* homologs in *A. thaliana*. C) Boxplots summarizing the fold change in root length of plant lines with SynTF-based activation of *DELLA* expression (orange) compared to a population of no gRNA control plant lines (gray, n=4). Every dot of the same color within each boxplot corresponds to an independent biological replicate of the same genotype. Different letters represent statistically significant differences (One-way ANOVA followed by Tukey HSD test, *p* < 0.05).

**Supplementary Figure S6.**
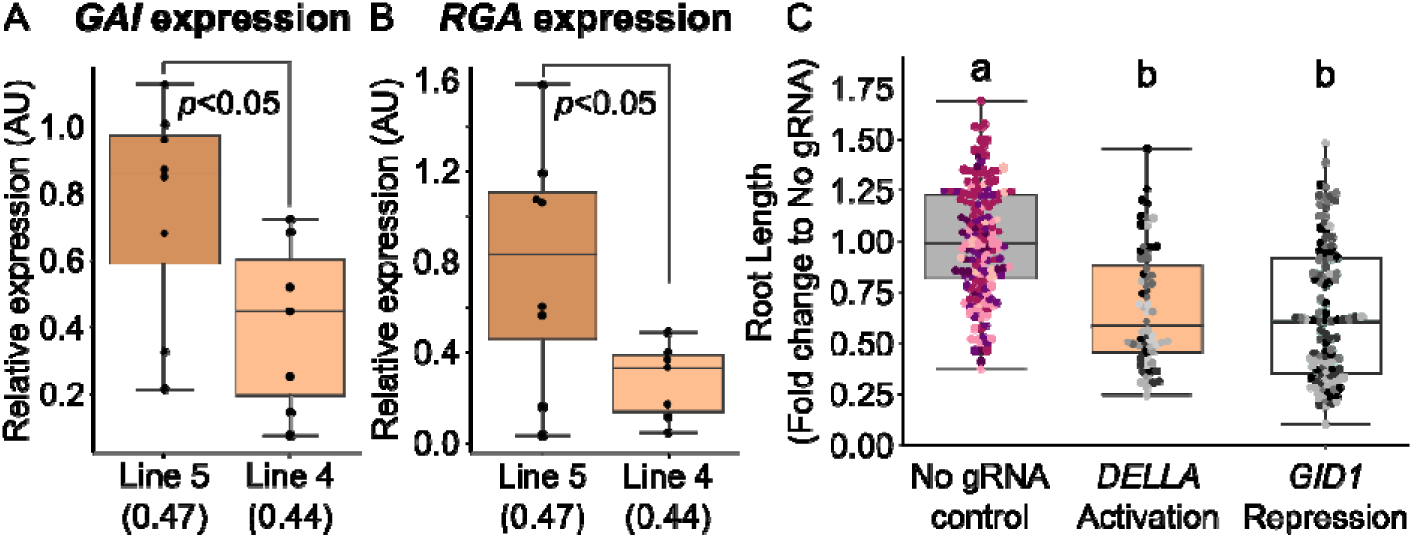
A,B) Boxplots summarizing significant differences in relative expression of both *GAI* (A) and *RGA* (B) in two *DELLA* activation lines, namely Line 5 (0.47-fold decrease in root length compared to no guide control, dark orange) and Line 4 (0.44-fold decrease in root length compared to no guide control, light orange). Each dot represents an independent biological replicate. C) Boxplots comparing the relative difference in fold change of root length across plant lines with either SynTF-based repression of *GID1* expression (teal, n=5) or SynTF-based activation of *DELLA* expression (orange, n=5) to a population of no gRNA control plant lines (gray, n=4). Different letters represent statistically significant differences (One-way ANOVA followed by Tukey HSD test, *p* < 0.05).

**Supplementary Figure S7.**
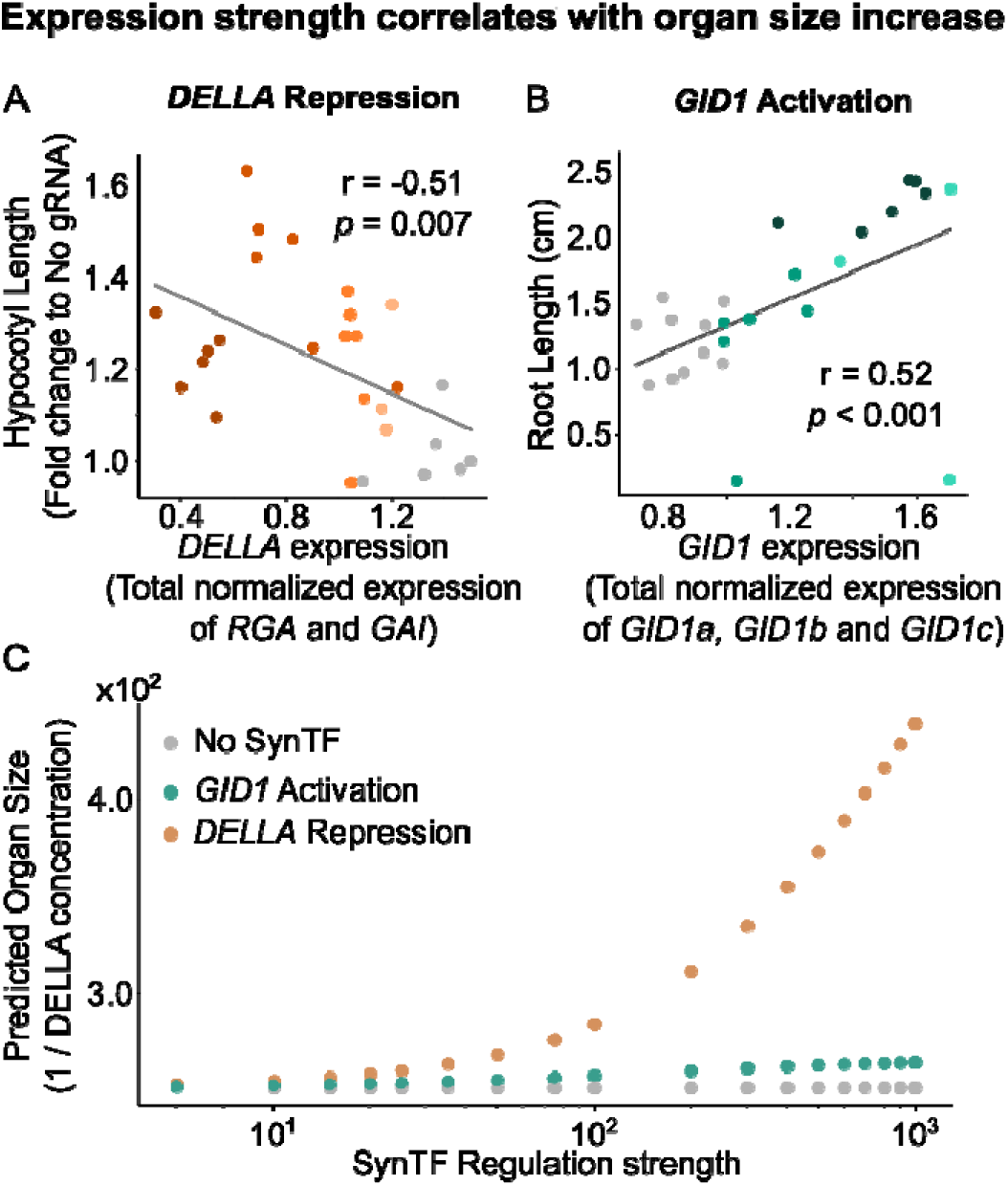
A) Scatterplot that demonstrates the Pearson’s correlation (r=-0.51, *p* = 0.007) between the fold change in hypocotyl length of the DELLA repression lines compared to the no gRNA controls and the sum of normalized *RGA* and *GAI* expression across four *DELLA* repression lines (progressively darker shades of orange) and the no gRNA control population (gray, n=1). Every dot of the same color corresponds to an independent biological replicate. B) Scatterplot that demonstrates the Pearson’s correlation (r=0.52, *p* < 0.001) between primary root length and the sum of normalized *GID1a*, *GID1b*, and *GID1c* expression across three *GID1* activation lines (progressively darker shades of teal) and the no gRNA control population (gray, n=3). Every dot of the same color corresponds to an independent biological replicate. C) Plot summarizing the predicted reciprocal of *DELLA* protein concentration at steady state, used as a proxy for organ size, from simulations of the GA signaling network with either no SynTF (gray), an activator targeted to regulate *GID1* expression (dark teal), or a repressor targeted to regulate *DELLA* expression (dark orange) across a range of different SynTF regulation strengths.

**Supplemental Figure S8.**
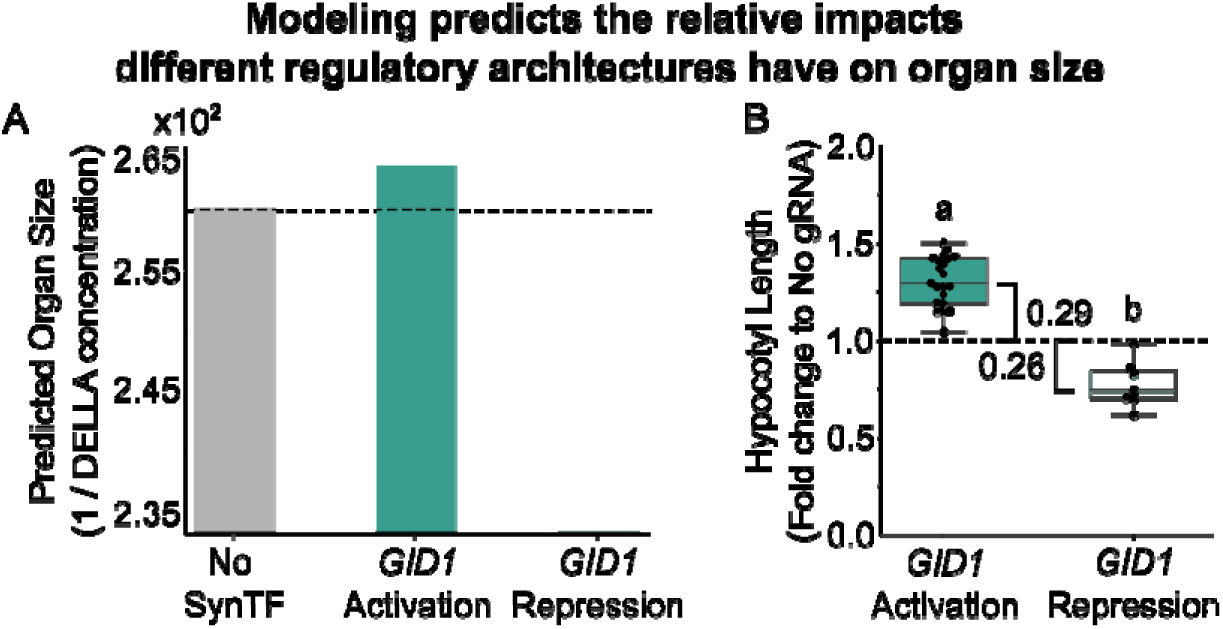
A) Bar plots summarizing the predicted reciprocal of DELLA protein concentration at steady state, used as a proxy for organ size, from simulations of the GA signaling network with either no SynTF (gray) and an activator (dark teal) or a repressor (teal) targeted to regulate *GID1* expression. F) Boxplots summarizing the fold change in hypocotyl length compared to the no gRNA controls of the *GID1* repression (teal) or *GID1* activation (dark teal) line with the strongest phenotype. The brackets represent the difference in fold change of hypocotyl length compared to the no gRNA controls. In all plots, the dashed black line highlights the phenotype of the no gRNA control. Letters represent statistically significant differences (One-way ANOVA followed by Tukey HSD test, *p* < 0.05)

**Supplemental Figure S9.**
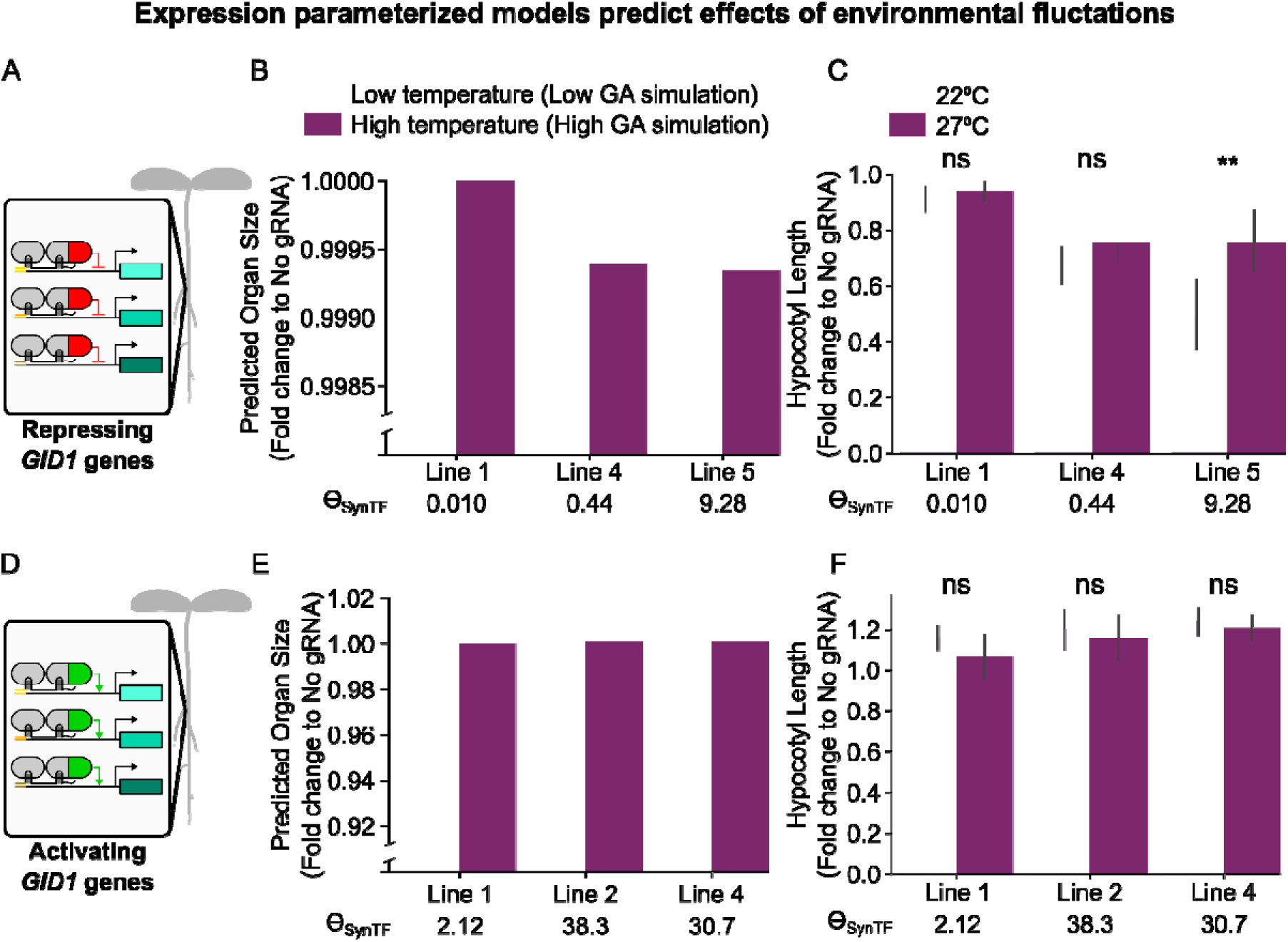
Expression parameterized models enable quantitative prediction of organ size. A) Schematic of the genetic circuit designed to implement SynTF-based repression on the three *GID1* paralogs in *A. thaliana*. B) Bar plots summarizing the predicted organ size, reported as the fold change in 1/DELLA concentration to the no gRNA control at low (light purple) and high (dark purple) GA simulations, to represent low and high temperatures, of the *GID1* repression lines. The values under the Lines on the x-axis represent their respective lt_SynTF_ value. C) Bar plots summarizing the fold change in hypocotyl length of the *GID1* repression lines compared to the no gRNA controls at low (light purple, 22⁰C) and high (dark purple, 27⁰C) temperatures. The values under the Lines on the x-axis represent their respective lt_SynTF_ value. D) Schematic of the genetic circuit designed to implement SynTF-based activation on the three *GID1* paralogs in *A. thaliana*. E) Bar plots summarizing the predicted organ size, reported as the fold change in 1/DELLA concentration to the no gRNA control at low (light purple) and high (dark purple) GA concentrations to represent low and high temperatures, of the *GID1* activation lines. The values under the Lines on the x-axis represent their respective lt_SynTF_ value. F) Bar plots summarizing the fold change in hypocotyl length of the *GID1* activation lines compared to the no gRNA controls at low (light purple, 22⁰C) and high (dark purple, 27⁰C) temperatures. The values under the Lines on the x-axis represent their respective lt_SynTF_ value. Statistical significance calculated using Welch’s two sample *t*-test (*p* < 0.05), * corresponds to *p* < 0.05, ** corresponds to *p* < 0.005, and *** corresponds to *p* < 0.0005.

**Supplementary Figure S10.**
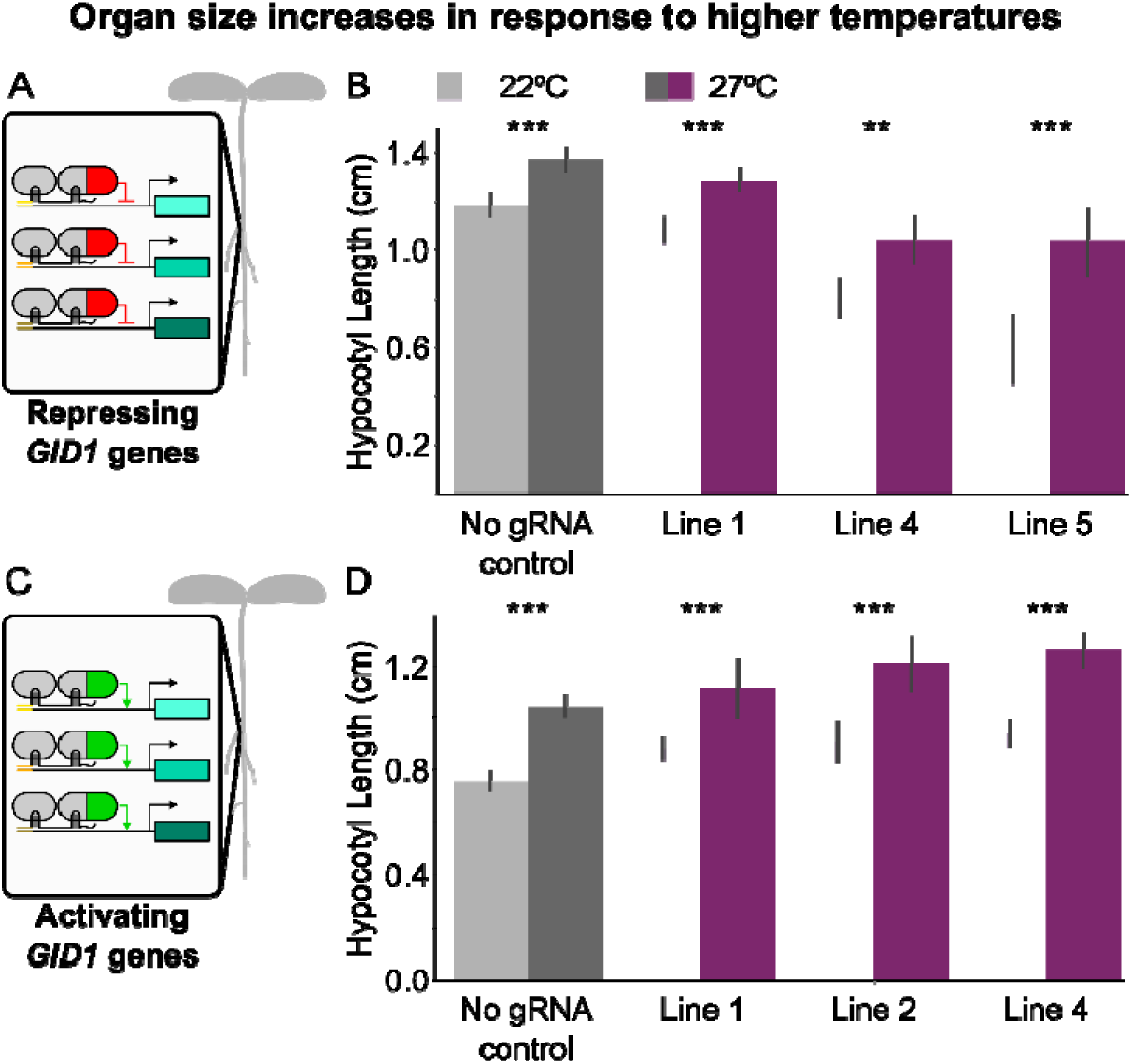
A) Schematic of the genetic circuit designed to implement SynTF-based repression on the three *GID1* paralogs in *A. thaliana.* B) Bar plots demonstrating hypocotyl length of eight-day old seedlings in either 22⁰C (light gray, light purple) or 27⁰C (dark gray, dark purple) of the no gRNA control population (gray) and the *GID1* repression lines (purple). C) Schematic of the genetic circuit designed to implement SynTF-based activation on the three *GID1* paralogs in *A. thaliana.* D) Bar plots demonstrating hypocotyl length of eight-day old seedlings in either 22⁰C (light gray, light purple) or 27⁰C (dark gray, dark purple) of the no gRNA control population (gray) and the *GID1* activation lines (purple). Statistical significance calculated using Welch’s two sample *t*-test (*p* < 0.05), * corresponds to *p* < 0.05, ** corresponds to *p* < 0.005, and *** corresponds to *p* < 0.0005.

**Supplementary Figure S11.**
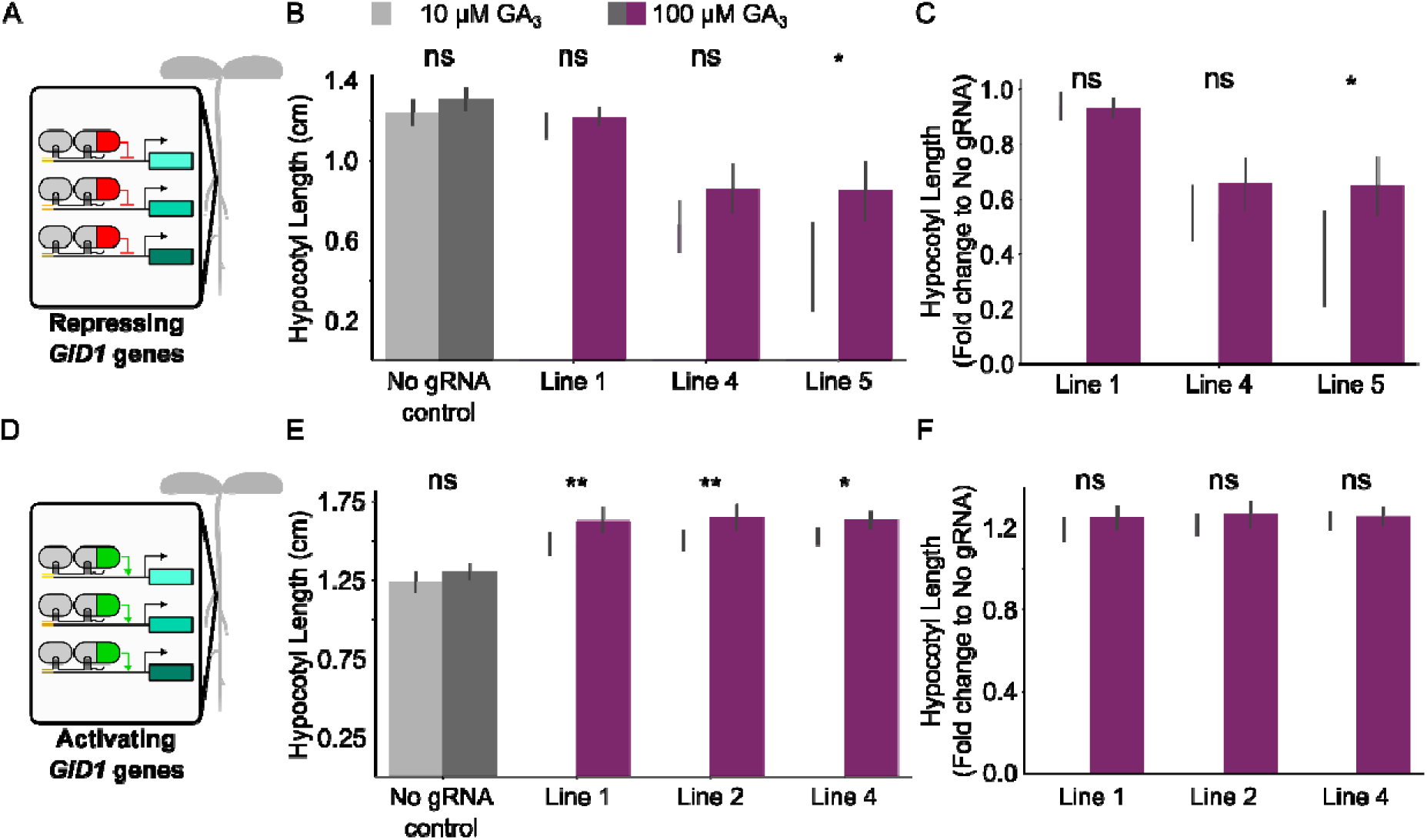
GA variations produce model predicted phenotypic effects on GID1 repression lines. A) Schematic of the genetic circuit designed to implement SynTF-based repression on the three *GID1* homologs in *A. thaliana.* B) Bar plots summarizing the hypocotyl length of the no gRNA controls (gray, n=3) and the *GID1* repression lines at low (light purple, 10 µM) and high (dark purple, 100 µM) GA_3_. C) Bar plots summarizing the fold change to no gRNA controls (gray, n=3) of the observed hypocotyl length phenotypes of three independent *GID1* repression lines supplemented with low (light purple, 10 µM) and high (dark purple, 100 µM) GA_3_. D) Schematic of the genetic circuit designed to implement SynTF-based activation on the three *GID1* homologs in *A. thaliana.* E) Bar plots summarizing the hypocotyl length of the no gRNA controls (gray, n=3) and the *GID1* activation lines at low (light purple, 10 µM) and high (dark purple, 100 µM) GA_3_. C) Bar plots summarizing the fold change to no gRNA controls (gray, n=3) of the observed hypocotyl length phenotypes of three independent *GID1* activation lines supplemented with low (light purple, 10 µM) and high (dark purple, 100 µM) GA_3_. Statistical significance calculated using Welch’s two sample *t*-test (*p* < 0.05), * corresponds to *p* < 0.05, ** corresponds to *p* < 0.005, and *** corresponds to *p* < 0.0005.

**Supplementary Figure S12.**
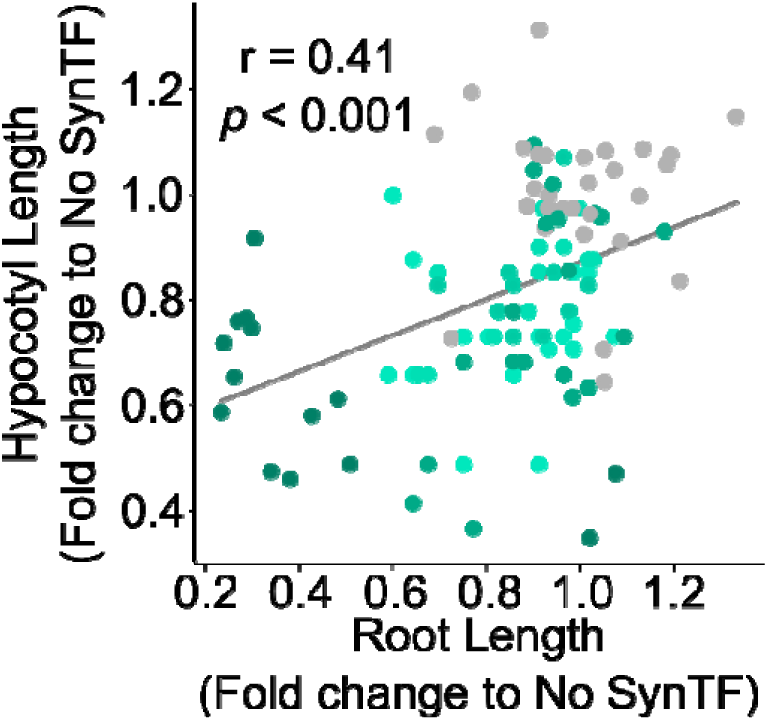
SynTF based repression of GID1s generates consistent dwarfing across tissues in tomato. Scatterplot summarizing the Pearson’s correlation between the fold change, compared to no gRNA control, in hypocotyl length and primary root length. Every dot of the same color corresponds to an independent biological replicate of the same genotype.

**Supplemental Table S1.** Summary of parameter values used in the full model.

| Parameter | Default Value | Meaning and Reference |
| --- | --- | --- |
| $l_a$ | $1.35 \mu\text{M}^{-1} \text{min}^{-1}$ | Association rate of GA<sub>4</sub> to GID1, (22) |
| $l_d$ | $2.84 \text{min}^{-1}$ | Dissociation rate of GA<sub>4</sub> to GID1, (22) |
| $p$ | $0.0776 \text{min}^{-1}$ | Rate of GID1 lid opening, (22) |
| $q$ | $0.0251 \text{min}^{-1}$ | Rate of GID1 lid closing, (22) |
| $u_{a1}$ | $10 \mu\text{M}^{-1} \text{min}^{-1}$ | GID1c.GA<sub>4</sub> association rate with DELLA binding site 1, (22) |
| $u_{a2}$ | $316.22 \mu\text{M}^{-1} \text{min}^{-1}$ | GID1c.GA<sub>4</sub> association rate with DELLA binding site 2, (22) |
| $u_{d1}$ | $0.133 \text{min}^{-1}$ | GID1c.GA<sub>4</sub> dissociation rate with DELLA binding site 1, (22) |
| $u_{d2}$ | $2.82 \text{min}^{-1}$ | GID1c.GA<sub>4</sub> dissociation rate with DELLA binding site 2, (22) |
| $u_m$ | $6.92 \text{min}^{-1}$ | Rate of tagging DELLA for degradation, from GID1c.GA<sub>4</sub> bound to DELLA binding site 1, (22) |
| $k_{a12}$ | 2904 | Association rate of GA<sub>12</sub> with GA20ox, (22) |
| $k_{a15}$ | 1072 | Association rate of GA<sub>15</sub> with GA20ox, (22) |
| $k_{a24}$ | 3099 | Association rate of GA<sub>24</sub> with GA20ox, (22) |
| $k_{a9}$ | 2073 | Association rate of GA<sub>9</sub> with GA3ox, (22) |
| $k_{d12}$ | 2.67 | Dissociation rate of GA<sub>12</sub> from GA20ox, (22) |
| $k_{d15}$ | 0.0088 | Dissociation rate of GA<sub>15</sub> from GA20ox, (22) |
| $k_{d24}$ | 0.0159 | Dissociation rate of GA<sub>24</sub> from GA20ox, (22) |
| $k_{d9}$ | 0.0088 | Dissociation rate of GA<sub>9</sub> with GA3ox, (22) |
| $k_{m12}$ | 198.8 | Rate of conversion of GA<sub>12</sub> to GA<sub>15</sub>, (22) |
| $k_{m15}$ | 763.8 | Rate of conversion of GA<sub>15</sub> to GA<sub>24</sub>, (22) |
| $k_{m24}$ | 2.6 | Rate of conversion of GA<sub>24</sub> to GA<sub>9</sub>, (22) |
| $k_{m9}$ | 763.8 | Rate of conversion of GA<sub>9</sub> to GA<sub>4</sub>, (22) |
| $\mu_{GA12}$ | $0.291 \text{min}^{-1}$ | GA degradation for *ga1-3* mutant and WT plants, (22) |
| $\omega_{GA12}$ | 0.0066 | Supply rate of GA12 in the *ga1-3* mutant, (22) |
| $\varphi_{GID1m}$ | $0.0457 \text{min}^{-1}$ | GID1 mRNA decay, (22) |
| $\varphi_{GA3oxm}$ | $0.103 \text{min}^{-1}$ | GA3ox mRNA decay, (22) |
| $\varphi_{GA20oxm}$ | $0.0478 \text{min}^{-1}$ | GA20ox mRNA decay, (22) |
| $\varphi_{DELLAm}$ | $0.0708 \text{min}^{-1}$ | DELLA mRNA decay, (22) |
| $\varphi_{SynTFm}$ | $0.0708 \text{min}^{-1}$ | SynTF mRNA decay, assumed to be the same as $\varphi_{DELLAm}$ |
| $\theta_{GIDm}$ | $5.6 \times 10^{-4}$ | DELLA protein for half maximal *GID1* transcription, (22) |
| $\theta_{GA3oxm}$ | 0.0082 | DELLA protein for half maximal GA3ox transcription, (22) |
| $\theta_{GA20oxm}$ | 0.638 | DELLA protein for half maximal GA20ox transcription, (22) |
| $\theta_{DELLAm}$ | 0.01 | DELLA protein for half maximal *DELLA* transcription, (22) |
| $\theta_{SynTFm}$ | 0.01 | DELLA protein for half maximal SynTF transcription, assumed to be the same as $\theta_{DELLAm}$ |
| $\delta_{GID1}$ | $19.3 \text{min}^{-1}$ | Translation rate, GID1 mRNA to GID1 protein, (22) |
| $\delta_{GA3ox}$ | $0.193 \text{min}^{-1}$ | Translation rate, GA20ox mRNA to GA3ox protein, (22) |
| $\delta_{GA20ox}$ | $0.193 \text{min}^{-1}$ | Translation rate, GA20ox mRNA to GA20ox protein, (22) |
| $\delta_{DELLA}$ | $5.28 \times 10^{-4} \text{min}^{-1}$ | Translation rate, DELLA mRNA to DELLA protein, (22) |
| $\delta_{SynTF}$ | $0.193 \text{ min}^{-1}$ | Translation rate, SynTF mRNA to SynTF protein, assumed to be the same as GA20ox |
| $\mu_{GID1}$ | $3.51 \text{ } \mu\text{M min}^{-1}$ | Rate of degradation of GID1 protein, (22) |
| $\mu_{GA3ox}$ | $3.51 \text{ } \mu\text{M min}^{-1}$ | Rate of degradation of GA3ox protein, (22) |
| $\mu_{GA20ox}$ | $3.51 \text{ } \mu\text{M min}^{-1}$ | Rate of degradation of GA20ox protein, (22) |
| $\mu_{DELLA}$ | $3.51 \text{ } \mu\text{M min}^{-1}$ | Rate of degradation of DELLA protein, (22) |
| $\mu_{SynTF}$ | $3.51 \text{ } \mu\text{M min}^{-1}$ | Rate of degradation of the SynTF protein, assumed to be the same as $\mu_{DELLA}$ |
| $\theta_{SynTF1}$ | 0 | Assumed, SynTF regulation strength for the no guide control population |
| $\theta_{SynTF2}$ | $9.931563 \times 10^{-3}$ | Fitted, SynTF regulation strength for *GID1* Repression Line 1 |
| $\theta_{SynTF3}$ | 0.04498460 | Fitted, SynTF regulation strength for *GID1* Repression Line 4 |
| $\theta_{SynTF4}$ | 9.277393 | Fitted, SynTF regulation strength for *GID1* Repression Line 5 |
| $\theta_{SynTF5}$ | $2.482093 \times 10^{-4}$ | Fitted, SynTF regulation strength for *GID1* Repression Line 2 |
| $\theta_{SynTF6}$ | 2.124854 | Fitted, SynTF regulation strength for *GID1* Activation Line 1 |
| $\theta_{SynTF7}$ | 38.27047 | Fitted, SynTF regulation strength for *GID1* Activation Line 2 |
| $\theta_{SynTF8}$ | 30.70163 | Fitted, SynTF regulation strength for *GID1* Activation Line 4 |

**Supplemental Table S2.** List of plasmids.

| Plasmid # | Plasmid Description | Link to Annotated Plasmid Maps |
| --- | --- | --- |
| P210 | pMOD_B_pGmUbi::NLS-MCP-DREB2A-tUBQ1<sup>7</sup> | https://benchling.com/s/seq-MwAvZNSQJKqmA7jjQk1k?m=slm-ZNVokLxETBFbsCeVualr |
| P218 | pMOD_C_p35s:NLS-PCP-TPLN300:tNos<sup>7</sup> | https://benchling.com/s/seq-oeZenFJYD4aXba0VMmo1?m=slm-LaJYSfMpSU2n9dshlt84 |
| P251 | pMOD_D-p35s:AtDELLA_guides1-1xMS2:t35s<sup>7</sup> | https://benchling.com/s/seq-34IAd2Ak7qPAOPtTvCAs?m=slm-w694bkGtyaDy3b5zhTb7 |
| P387 | pMOD_A_pUBQ10:NLS-Cas9-tHSP<sup>7</sup> | https://benchling.com/s/seq-gcG6pGbgeOKdE49GSFEO?m=slm-TdLwqOoOyOaqvvucPXE7 |
| P671 | pTRANS_220d-pUBQ10:NLS-AtCas9:tHSP - pGmUbi:NLS-MCP-DREB2A-tUBQ1 - p35s:NLS-PCP-TPLN300:tNos - p35s:AtDELLA_guides1-1xMS2:t35s | https://benchling.com/s/seq-mma2v82CeUQxRfDbdVF?m=slm-KRvmmsgJFG25S8quBPrvg |
| P683 | pTRANS_220d-pUBQ10:NLS-AtCas9:tHSP - pGmUbi:NLS-MCP-DREB2A-tUBQ1 - p35s:NLS-PCP-TPLN300:tNos - p35s:AtGID_guides1-1xPP7:t35s | https://benchling.com/s/seq-qKv521SztV10SKalMc3t?m=slm-oeJhMAjbWFHnsnHL3gZM |
| P684 | pTRANS_220d-pUBQ10:NLS-AtCas9:tHSP - pGmUbi:NLS-MCP-DREB2A-tUBQ1 - p35s:NLS-PCP-TPLN300:tNos - p35s:AtGID_guides1-2xPP7:t35s | https://benchling.com/s/seq-vYJisQVdibhGCT1twsBJ?m=slm-ntsZ2SOAqxoaHdDZtp3M |
| P710 | pMOD_D-p35s:DELLA-guides-2xPP7-t35s<sup>7</sup> | https://benchling.com/s/seq-25g28HyVAIDdktL1qLL6?m=slm-BhxHy2z7olobWEfcylLU |
| P731 | pMOD_D-p35s:GID-guides-1xMS2-t35s<sup>7</sup> | https://benchling.com/s/seq-e67WY9o1PfYe5VvBaZ1F?m=slm-z2sjl8sWoZKQMWHFALMf |
| P735<sup>7</sup> | pMOD_D1'-p35s:AtGID_guides1-1xPP7<sup>7</sup> | https://benchling.com/s/seq-qU5SNDnPoYaabhH75pW3?m=slm-b9VXW7CuCIAoAtOI9HHz |
| P768<sup>7</sup> | pMOD_D1'-p35s:AtGID_guides1-2xPP7<sup>7</sup> | https://benchling.com/s/seq-l7vXkJhLRhQarZoLSi27?m=slm-5Q9ARa1MZtOApbqkLY9Z |
| P860<sup>7</sup> | pMOD_D2'-p35s:AtGID_guides2-1xPP7<sup>7</sup> | https://benchling.com/s/seq-cgas2gwlwy4O9YwY30LI?m=slm-DS2ZnZas1snmlmWMcFVU |
| P861<sup>7</sup> | pMOD_D2'-p35s:AtGID_guides2-2xPP7<sup>7</sup> | https://benchling.com/s/seq-4dU0OooeiqPd5ZYSpy3H?m=slm-Q5io5eJT02jhiNgEgPAY |
| P925 | pTRANS_220d-pUBQ10:NLS-Cas9-tHSP- | https://benchling.com/s/seq- |
|  | pGmUbi:NLS-MCP-DREB2A-tUBQ1-p35s:NLS-PCP-TPLN300:tNos-p35s:GID-guides1-2xPP7-t35s - p35s:GID-guides2-2xPP7-t35s | [rbcGB2xbNlubWVsh031c?m=slm-SgAUseEU9cvW8lpwaW2Z](https://benchling.com/s/seq-rbcGB2xbNlubWVsh031c?m=slm-SgAUseEU9cvW8lpwaW2Z) |
| P1212 | pTRANS_220d-pUBQ10:NLS-Cas9-tHSP-pGmUbi:NLS-MCP-DREB2A-tUBQ1-NLS-PCP-TPLN300:tNos-p35s:DELLA-guides-2xPP7-t35s | https://benchling.com/s/seq-Szsa9qRgv2rZRLKGh51F?m=slm-2bd96Ox94IZwMRabNePn |
| P1317 | pTRANS_220d-pUBQ10:NLS-Cas9-tHSP-pGmUbi:NLS-MCP-DREB2A-tUBQ1-p35s:NLS-PCP-TPLN300:tNos-p35s:GID-guides-1xMS2-t35s | https://benchling.com/s/seq-Obu7yyCpOC9QV0nPTKPB?m=slm-5hF50ttCvqTvwuyMuBnI |

**Supplemental Table S3.** List of sequences targeted by sgRNAs.

| **Guide name** | **Sequence (3' to 5')** |
| --- | --- |
| GID1a_Target1 | AGGGATGAGTAGGG |
| GID1a_Target2 | TGTTGGGTCATGGG |
| GID1a_Target3 | TGCGCGCATAAGTT |
| GID1a_Target5 | GGATAACAACAAA |
| GID1a_Target6 | GTCAATTTTATTCG |
| GID1b_Target1 | GACCAATCGGACGG |
| GID1b_Target2 | TACACATGACACCG |
| GID1b_Target3 | AGCACTAATGAGAG |
| GID1b_Target5 | GAATTAGGAAGCAG |
| GID1b_Target6 | GAATTAGGAAGCAG |
| GID1c_Target1 | AAGAATATCGGCGT |
| GID1c_Target2 | CGACTGTTGTTGGG |
| GID1c_Target3 | TGCGAACAGATTAA |
| GID1c_Target5 | TCAGAATACTAGAA |
| GID1c_Target6 | CACCCTTGGGAGCA |
| RGA_Target1 | CATCTAAAGTGATA |
| GAI_Target1 | TGTCATGCAACTAG |
| RGL1_Target1 | TTGCGGTTGTACCA |
| RGL2_Target1 | TTCTTAAACCCAAA |
| RGL3_Target1 | CGGGCTGTTACTTT |

**Supplemental Table S4.**
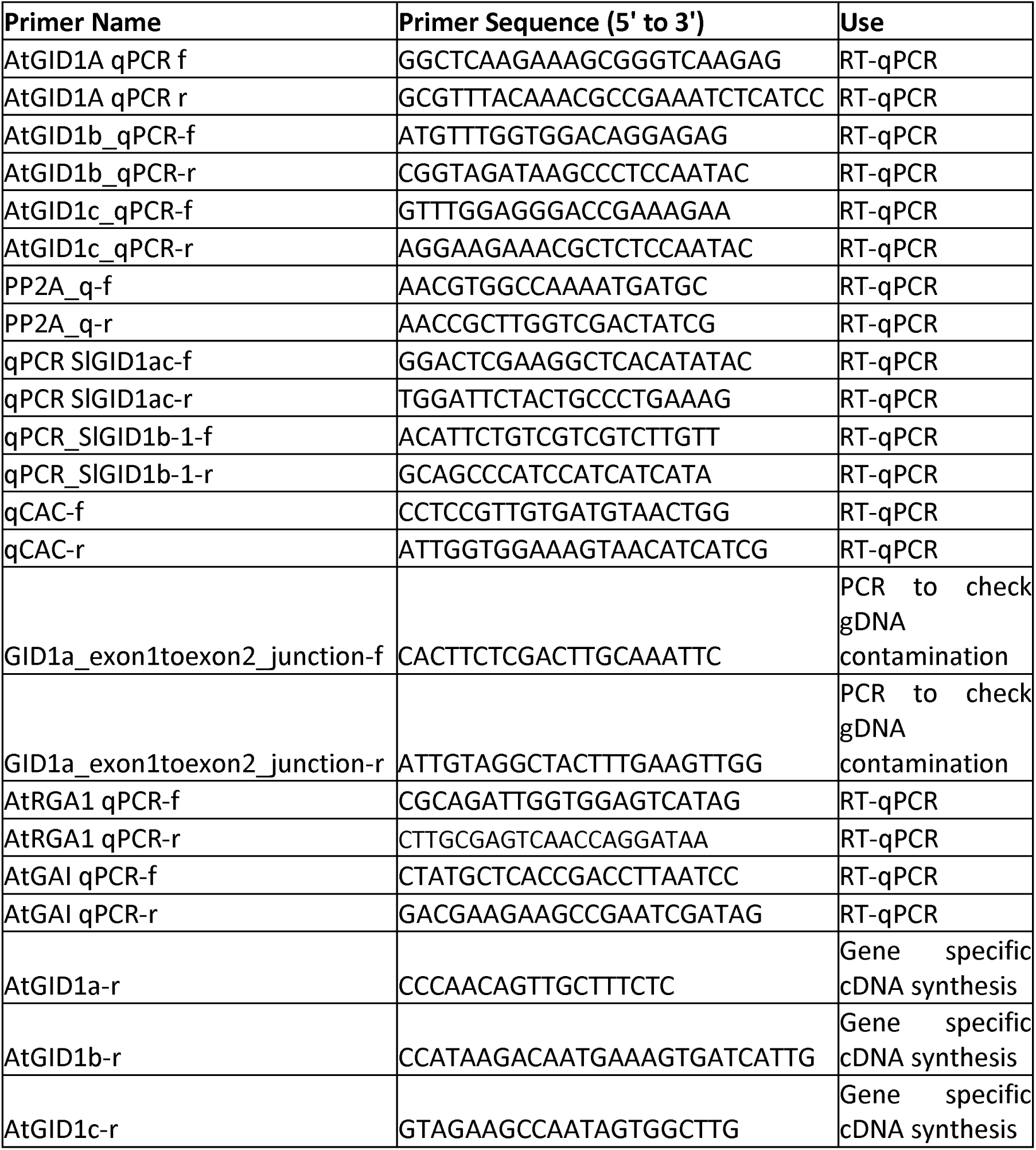
Gene accessions referenced in the manuscript.

**Supplemental Table S5.**
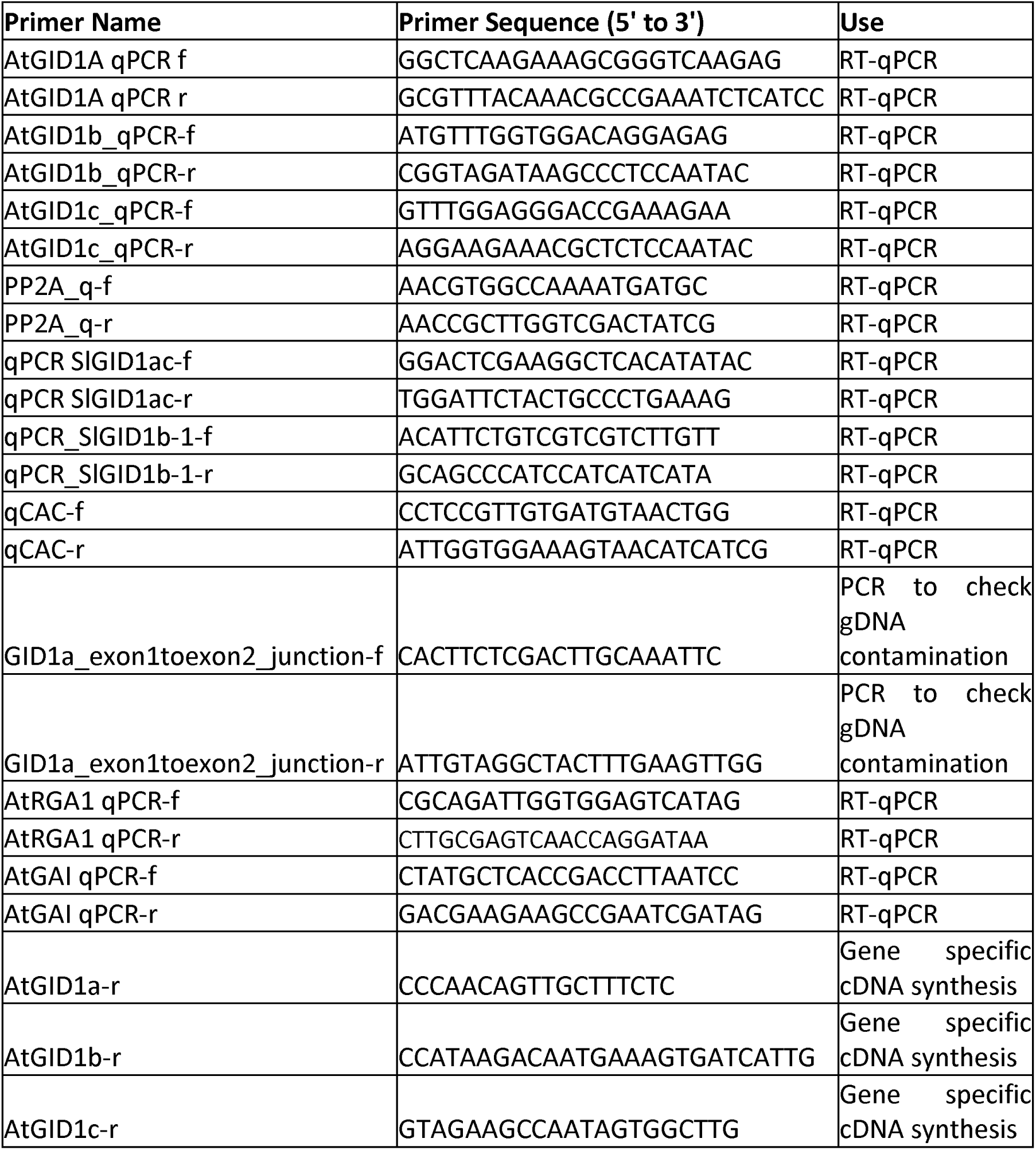
List of primers used in this manuscript.

## Supplemental Materials and Methods

### Model simulations

A modified version of the mathematical model of GA signaling, originally developed by Middleton et al. (35), was implemented in python to simulate the impact of SynTF-based regulation of *GID1* or *DELLA* expression on steady state *DELLA* protein levels. This involved adding equations to simulate SynTF transcription. (Eq. 1) and translation (Eq. 2) as well as modifications to the Hill equations simulating *GID1* activation (Eq. 3) or repression (Eq. 4), or *DELLA* activation (Eq. 5) or repression (Eq. 6). These are represented in **Fig. S1** and shown below:

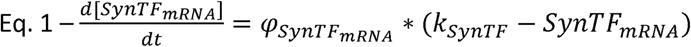

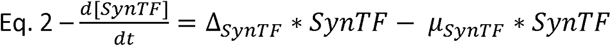

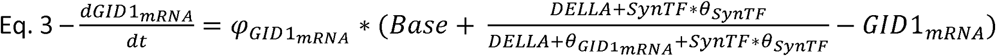

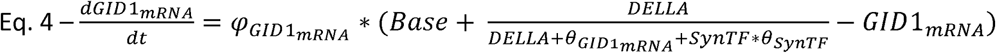

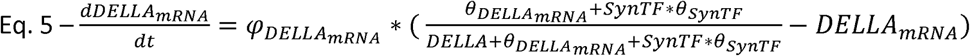

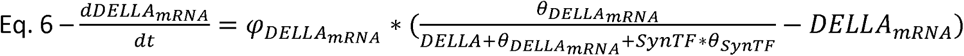

The constants for these equations (<p-scaler for transcription, Δ-rate of translation, µ-rate of protein decay, *Base*-basal rate of *GID1* expression) were either kept the same as the original Middleton *et al*. model or were assumed to be the same as the DELLA for those associated with the SynTF (**Table S1**). The values used are detailed in the associated jupyter notebook on github (https://github.com/arjunkhakhar/250610_GA_signaling_paper). For the SynTF, we explored a broad range of both expression levels (k_SynTF_) and regulation strengths (⍰_SynTF_). To analyze steady state DELLA concentrations when SynTFs are targeted to regulate either *GID1* or *DELLA*, we picked initial conditions where all variables were set to zero and ran a numerical solver (odeint) from the scipy package with enough timesteps for all variables to be at, or asymptotically approach, steady state. The inverse of steady state DELLA concentrations, which we assume to be a proxy for organ size, were then plotted using functions from the matplotlib and seaborn packages.

We also explored whether this model could be used to predict how the strength of GID1 repression would alter organ size, specifically root length as this is what the model parameters were based on, relative to the no gRNA plant. We did this by first using the measured *GID1* repression levels to calculate a ⍰_SynTF_ for each line, predicting organ size for each line using this ⍰_SynTF_, and finally comparing the measured fold change in root length to this prediction. More specifically, we first used the model of GA signaling with *GID1* repression (Eq. 4) with the ⍰_SynTF_ set to 0 to simulate the no gRNA control plant. Using this we calculated the GID1 mRNA concentration at steady state. We then used the fold change in expression of *GID1* in each *GID1* repression line compared to the no gRNA control population, measured via RT-qPCR, to determine the target GID1 mRNA concentration we would expect in that line. This is then used to determine the ⍰_SynTF_ that would generate this level of GID1 mRNA at steady state. Next, we simulated what the DELLA protein concentration would be at steady state for each *GID1* repression line’s ⍰_SynTF_. We then compared the measured fold change in root length in these lines to the fold change in predicted 1/DELLA protein concentration scaled to be between the same range as root length measurements.

Finally, we explored the impact of environmentally generated fluctuations in GA levels on the phenotypes generated by modulation of GA signaling using SynTFs. To do this we built modified versions of the model where the concentration of GA_4_, bioactive GA, was set to a constant level and then the steady state DELLA concentration was simulated. This was performed for each line’s ⍰_SynTF_. As part of this analysis we also calculated ⍰_SynTF_ values for three *GID1* activation lines using the same approach as detailed above. As prior literature demonstrates high temperatures lead to higher GA in plant tissues (41), we simulated and plotted 1/DELLA at steady state for GA_4_ set to 350 and 500 representing normal and high temperatures, respectively. The code for this can be found on github (https://github.com/arjunkhakhar/250610_GA_signaling_paper).

### Plasmid construction

All plasmids constructed in this work were built using Golden Gate Assembly and Modular Cloning (47, 48) and are listed with links to their annotated plasmid maps in **Supplemental Table S2**. Codon optimized sequences of nuclease active Cas9, PCP, MCP, and TPLN300 with correct restriction site overhangs were amplified from previously reported plasmids (11). The sequence for the DREB2A activation domain with restriction site overhangs was synthesized using Twist Bioscience. All sgRNA target sequences in this work were designed using CHOPCHOP v3 (49) and the target sequences are listed in **Supplemental Table S3** and the target gene accession numbers are listed in **Supplemental Table S4**. The gRNA sequences were added to various DNA parts as overhangs using primers and PCR and were reconstituted during the assembly and ligation process. Recruitment motif sequences, PP7 and MS2, used in this study were previously reported (11, 50). All plasmids used in this work will be available via AddGene.

### *Arabidopsis* root phenotyping

Seeds were surface sterilized in a solution containing 70% Ethanol and 0.05% Triton-X-100 for 20 minutes. Seeds were immediately rinsed with 95% Ethanol. Seeds were plated on 1/2x MS (52) containing 0.8% PhytoAgar™ (plantmedia Cat. No. 40100072-1) and Kanamycin (50mg/L). After plating the seeds, plates were stratified in the dark at 4⁰C for five days. After five days, plates were grown vertically under short day conditions (9hr light/15hr dark) at 22⁰C in a Percival growth chamber for 8 days. Plates were scanned using an Epson scanner and root length was measured using ImageJ.

### Gene expression analysis via RT-qPCR in *Arabidopsis*

For characterization of *GID1* and *DELLA* expression, whole eight-day-old seedlings from the root phenotyping assays of the different regulatory architectures and no gRNA control lines were harvested either during the day or one hour after dark and immediately frozen in liquid nitrogen. RNA was extracted using the QIAGEN RNeasy Plant Mini Kit (Cat No. 74904). The RNA samples were treated with the TURBO DNA-*Free^TM^*kit (Invitrogen Cat No. AM1907) to remove genomic DNA. Complementary DNA (cDNA) was synthesized using the LunaScript® RT SuperMix Kit or the LunaScript® RT primer free MasterMix Kit and gene specific primers. Proper removal of the genomic DNA was confirmed with PCR using GoTaq® Green Master Mix (Promega Cat No. M7122) and primers spanning the first and second intron of *GID1a*. Concentrations of *GID1a*, *GID1b*, *GID1c*, *RGA*, *GAI*, and *PP2A* were quantified using RT-qPCR performed with the Luna® Universal qPCR Master Mix on the BioRad CFX Opus qPCR machine. Cycle threshold values for *GID1* and *DELLA* were normalized to the housekeeping gene (*PP2A*). Two technical replicates were performed for all samples. The gene specific, PCR and RT-qPCR primers used are listed in **Supplemental Table S5**.

### *Arabidopsis* shoot phenotyping

For hypocotyl measurements, seeds of *A. thaliana* transgenic lines were surface sterilized and plated on media plates as described above. For the experiments investigating GA levels on organ size, GA_3_ was added to the media at concentrations of 10 and 100 µM. Plates were stratified for three days in the dark at 4⁰C, light shocked for six hours in a Percival growth chamber set to 22⁰C or 27⁰C (for the temperature treatments) and short-day conditions (9hr light/ 15hr dark) and then were wrapped in foil and placed vertically in the correct growth chamber to promote etiolation. After three days, plates were scanned using an Epson scanner and hypocotyls were measured using Image J.

For rosette and leaf phenotyping, seeds of *A*. *thaliana* transgenic lines were surface sterilized and plated as described above. Seeds were stratified for three days in the dark at 4⁰C, light shocked for six hours in a Percival growth chamber set to 22⁰C and long-day conditions (16hr light/8hr dark) and then were wrapped in foil and returned to the growth chamber. After three days, the foil was removed, and plates were returned to the growth chamber. After 15 days, 12 seedlings were transplanted into soil (2:1 Pro-Mix HP to Vermiculite mix) in 2.5-inch square pots and placed in a Conviron growth chamber set to 22⁰C, long-day conditions (16hr light/8hr dark), 150uM light intensity, and 60% relative humidity. Plants were placed in flats in a randomized block design and were bottom-watered every two to three days with tap water. After 15 days in the growth chamber, each plant was dissected by leaf and scanned in order of developmental age. Individual leaf areas were measured using ImageJ. Rosette areas were calculated by summing the areas of individual leaves for each plant.

For inflorescence phenotyping, seeds of *A. thaliana* transgenic lines were surface sterilized and plated on 1/2x MS media plates as described above. Plates were stratified for three days in the dark at 4⁰C and then transferred into Percival growth chamber set to 22⁰C and short-day conditions (8hr light/16hr dark). Three-week-old seedlings were transplanted into soil (2:1 Pro-Mix HP to Vermiculite mix) into 2.5-inch square pots. Plants were organized in a randomized block design in a greenhouse and bottom-watered by hand every two to three days. Plants were monitored daily and the date that individual plants transitioned to reproductive growth was collected. Once all plants had inflorescences that reached at least approximately two centimeters in height, inflorescence height was collected. Inflorescence height was normalized by days-post bolting for better comparisons of dwarfing between genotypes.

### Tomato phenotyping and gene expression analysis

Seeds of M82 WT, M82 Cas9, and *GID1* repression tomato lines were surface sterilized in 10mL of 20% commercial bleach with two drops of Tween20 with constant agitation for 20 minutes. Seeds were rinsed three times with sterile water. Approximately 20 seeds per line were sown in Magenta boxes containing 1/2x MS supplemented with 50ug/mL Kanamycin and placed in a Percival growth chamber set to 25⁰C and long day (16hr light/8hr dark) conditions. Hypocotyl and root length of 14-day old seedlings were measured and seedlings were transplanted into ProMix HP soil in 4-inch square pots and placed in a greenhouse with no active temperature or light control. After six days and 34 days, epicotyl and stem lengths were measured, respectively.

To measure *GID1* expression in tomato plants, tissue was collected one hour after dark by sampling 14-day-old seedlings. Seedlings were dissected into hypocotyls and roots and immediately frozen in liquid nitrogen. Seedling tissue was stored at –80⁰C prior to lysing. Cells were lysed using a mechanical tissue lyser and RNA was extracted using TRIzol (53) or the QIAGEN RNeasy Plant Mini Kit (Cat No. 74904) followed by treatment with the TURBO DNA-*Free^TM^* kit (Invitrogen Cat No. AM1907) to remove genomic DNA. The LunaScript® RT SuperMix Kit was used to synthesize cDNA. Concentrations of *GID1a*, *GID1b-1*, *GID1b-2,* and *CAC* were quantified using RT-qPCR performed with the Luna® Universal qPCR Master Mix on the BioRad CFX Opus qPCR machine. Two technical replicates were performed for all samples. Lack of genomic DNA contamination was validated using an RT negative control during the RT-qPCR reaction. The PCR and RT-qPCR primers used are listed in **Supplemental Table S5**. Cycle threshold values of the *GID1* genes were normalized to the housekeeping gene (*CAC*).

